# Multivalent 28S rRNA Is the Organizer of the Nucleolus’s Multi-layered Architecture

**DOI:** 10.1101/2024.03.13.584914

**Authors:** Jieli Wei, Yize Zhang, Bingcheng Ye, Xueyuan Fan, Yucen Hu, ShengQi Xiang, Weirui Ma

## Abstract

The nucleolus widely exists in all eukaryotic species. Throughout evolution, two types of nucleoli have emerged: bipartite, which has two-layered subcompartments, and tripartite, which features three nested sub-compartments: granular component (GC), dense fibrillar component (DFC), and fibrillar center (FC). FC and DFC form a core-shell architecture and are immersed by a large GC. However, factors that mediated the formation of the three-layered architecture remain largely unknown. Here, we discovered that rRNAs have specific localizations within the nucleolus, with 28S rRNA forming assemblies surrounding the DFC. Multivalent interactions of 28S rRNA drove the formation of the hollow shell structure of DFC, which mediated the development of a three-layered structure resembling nucleolar GC-DFC-FC. Moreover, in organisms with a tripartite nucleolus, 28S rRNA has evolved to acquire a higher degree of multivalency. Our data implies that the emergence of multivalency of 28S rRNA facilitates the transition of bipartite to tripartite nucleolus.

## Introduction

The nucleolus is a highly distinguishable membrane-less organelle found extensively in eukaryotic cells. It plays roles in various fundamental biological processes^1,2^, for instance, rRNA synthesis and processing^1,2^, ribosome biogenesis^1,2^, assembly of RNPs (ribonucleoproteins)^3,4^, 3D genome folding^5,6^, and protein quality control^7,8^. Across the eukaryotic cells, there are two distinct types of nucleoli^9–11^: the tripartite nucleolus in baker’s yeast^12,13^ and amniotes^10^, including mammalians^1^, birds^10^, and lizards^11^, and the bipartite nucleolus in other eukaryotes^9–11^, including Drosophila^14^ and C.elegans^15^. the tripartite nucleolus comprises three internal subcompartments, layered from outside to inside: the granular component (GC), the dense fibrillar component (DFC), and the fibrillar center (FC) (Figure 1A). Under electron microscopy and super-resolution fluorescence microscopy^16,17^, the DFC exhibits a ring-like morphology, surrounding the spherical FC condensate, indicating the FC and DFC form a core-shell architecture. FC-DFC units are immersed by a large GC^1,16,17^. In contrast, the bipartite nucleolus consists of only two layers: the FC and GC^11^ (Figure 1B). Previous research has suggested that the emergence of the DFC occurs in the reptilia^9,11^ and is correlated with the expansion of sequence in the intergenic space between rDNA repeats^9^. However, the factors driving the transition from a two-layered to a three-layered nucleolus remain unclear.

**Figure 1.**
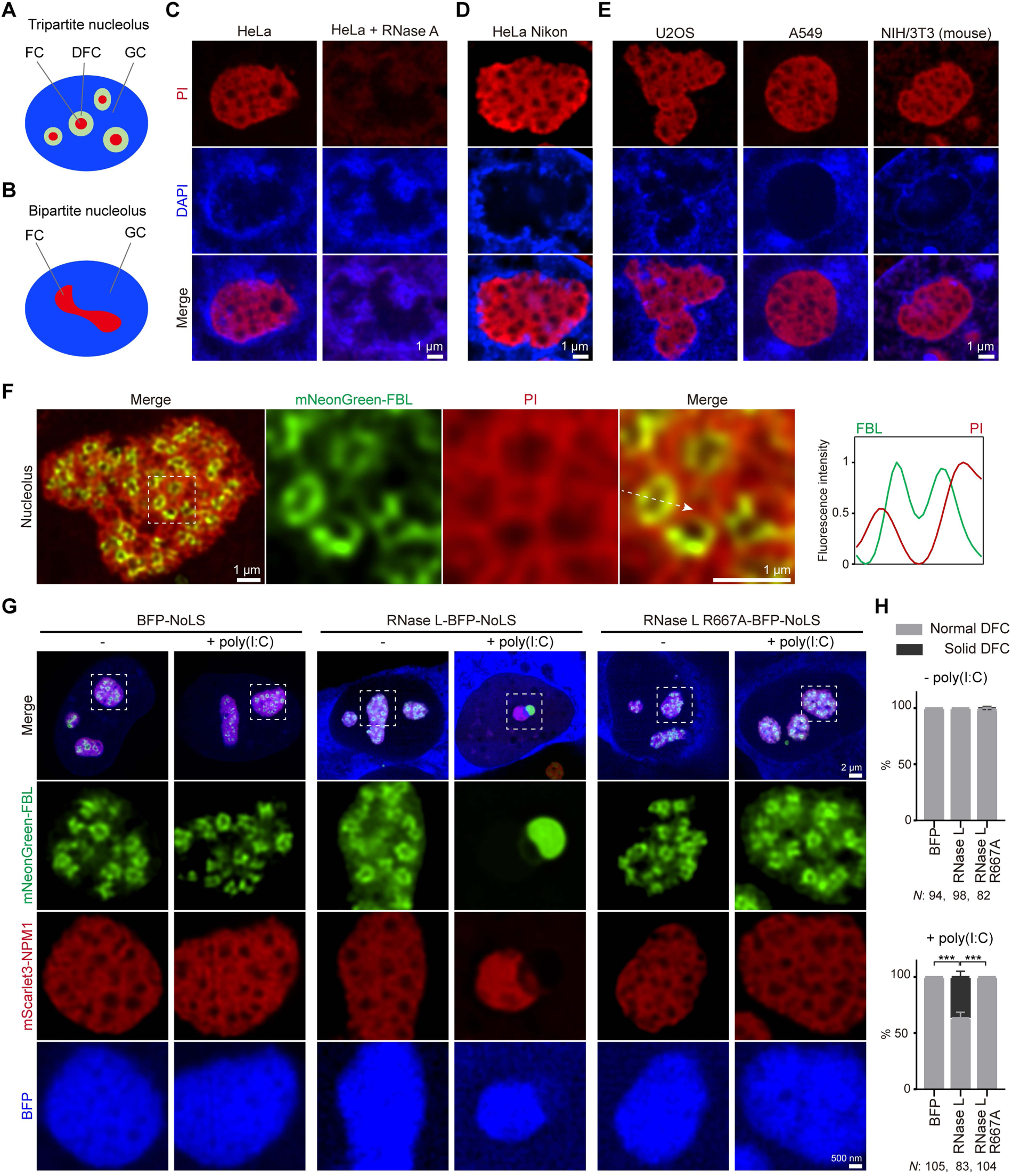
RNA is essential for the formation of the hollow shell architecture of DFC in cells. (**A**) Schematic of the tripartite nucleolus. GC, granular component; DFC, dense fibrillar component; FC, fibrillar center. (**B**) Schematic of the bipartite nucleolus. (**C**) Representative super-resolution images (Spin-SR with SoRa) of nucleoli in HeLa cells stained with PI and DAPI. (**D**) As in (C), but the images were taken using NIKON Confocal AX with NSPARC. (**E**) As in (C). but in U2OS, A549, and NIH/3T3 cells. (**F**) Representative super-resolution images of the nucleolar DFC and RNA (PI staining) in HeLa cells. Stably expressed mNeonGreen-FBL labels the DFC. Shown are two different magnifications. Right: line profile of fluorescence intensities. The arrow indicates the plane used for the line profile generation. (**G**) Fluorescent confocal microscopy of the nucleolus in HeLa cells before and after poly(I:C) transfection. mNeonGreen-FBL and mScarlet-NPM1 were stably expressed to label the nucleolar DFC and GC respectively. Prior to poly(I:C) transfection, BFP-NoLS, RNase L-BFP-NoLS, and RNase L R667A-BFP-NoLS were transiently transfected into HeLa cells. Bottom: Higher magnification of indicated regions (dotted boxes). (**H**) Quantification of the percent of cells with normal DFC or solid DFC in (**G**). Shown is mean ± SD of three biological replicates. *N*, the total number of cells transfected with indicated BFP-tagged proteins in each condition from three independent experiments. t test ,***p < 0.001.

The primary function of nucleoli is ribosomal biogenesis^1,2^, starting with rRNA transcription at the border of the FC^16^, where RNA polymerase I (Pol I) is enriched^17^. The rRNA is then transferred to the DFC for processing and subsequently to the GC for ribosomal subunit assembly^1^. It has been shown that DFC of an aberrant size is associated with a reduced level of rRNA transcription^18^. Additionally, the nucleolus exhibits diverse dynamics in its sub-organization^19^ during different developmental stages^20^, under cellular stresses^21^, during viral infections^22^, and in human diseases^23,24^, indicating a close link between sub-nucleolar organization and function^1^. Studying the formation of multi-layered nucleolar organization is valuable for understanding nucleolar functions.

Recent studies have demonstrated that the nucleolus forms through liquid-liquid phase separation (LLPS)^1,25–27^, with its multi-layered subcompartments representing immiscible liquid phases^1,28^. In vitro nucleolar reconstitution has shown that NPM1, a GC-localized protein, and FBL, a DFC-localized protein, form condensate through LLPS. The difference in surface tension between NPM1 and FBL condensates prevents the mixing of the two phases, thus allowing the formation of a two-layered, nucleolar-like architecture^28^. However, in this reconstitution, FBL forms solid spherical condensates^28^, unlike in the three-layered mammalian nucleoli, where the DFC appears as a hollow shell enclosing the FC. To date, the factors determining the formation of the DFC’s hollow shell structure remain unclear. In this study, we set out to identify these factors.

RNAs are critical regulators of LLPS^29^. RNA can phase separate on its own^30^ and promote the phase separation of proteins^29,31^, thereby nucleating the formation of membrane-less organelles, including paraspeckles^32^. RNA can also influence the biophysical properties^33,34^ and act as the skeleton of condensates, determining the three-dimensional morphology of RNA granules^35^. As the nucleolus contains numerous RNAs, we hypothesize that RNA is responsible for forming hollow shell structures of DFC.

## Results

### RNA forms assemblies surrounding the DFC in the nucleolus

To investigate the role of RNA in forming sub-nucleolar structures, we first examined the localization of RNAs in the nucleolus using propidium iodide (PI) and DAPI staining. PI, which stains both DNA and RNA, displayed strong fluorescence signals in the cytoplasm (Figure S1A) and the nucleolus (Figures 1C and S1A). Notably, in the nucleolus, the fluorescence from the DNA-specific dye DAPI was weaker than in the nucleoplasm. Furthermore, after treatment with RNase A, the PI fluorescence signal in both the cytoplasm (Figure S1A) and nucleolus was significantly reduced (Figures 1C and S1A), indicating that the nucleolar PI signal is primarily derived from RNA. Therefore, PI staining is indicative of RNA locations.

We then performed fluorescence imaging with two different super-resolution microscopy in various human and mouse cell lines (Figures 1C-1E). We observed that RNA within the nucleolus was not homogeneously distributed; instead, it exhibited a mesh-like morphology with multiple holes where the PI fluorescence signal was notably weaker (Figures 1C-1E). Given that PI is an intercalator binding to double-stranded DNA (dsDNA) and double-stranded RNA (dsRNA) by intercalating between bases, our findings suggest that RNA within the mammalian nucleolus forms conserved assemblies characterized by extensive RNA-RNA interactions.

We next labeled the nucleolar DFC using stably expressed mNeonGreen-FBL and performed PI staining in three human cell lines. In line with previous findings, FBL displayed a ring-like morphology under super-resolution microscopy (Figure 1F), indicative of the hollow shell architecture of DFC. Interestingly, the FBL rings were frequently situated within the holes of the RNA assembly (Figures 1F and S2A). The PI fluorescence signal was observed to encircle and partially overlap with the FBL rings. However, the signal was decreased at the center of the ring, which corresponds to the location of the fibrillar center (FC) (Figures 1F and S2A). This evidence indicates the close association between RNA assemblies and the multi-layered structures within the nucleolus.

### RNA is essential for the formation of the hollow shell architecture of DFC in cells

To investigate RNA’s involvement in the formation of the DFC’s shell structure, we employed an RNase L experiment to degrade RNA within the nucleolus in an inducible manner^36^. RNase L is an endoribonuclease activated in response to dsRNA virus infection in cells^37^. To devoid the potential interference of endogenous RNase L, we first created RNase L knockout cells using CRISPR/Cas9 technology (Figures S3A and S3B) and then generated stable cell lines expressing mNeonGreen-FBL and mScartlet3-NPM1 under the RNase L knockout background to label the nucleolar DFC and GC, respectively. Next, we transiently transfected these cells with RNase L-BFP, fused to a NoLS (nucleolus localization signal) to target a portion of RNase L to the nucleolus (Figure S4A). Subsequently, we transfected the cells with Poly(I:C), an analog of dsRNA, to activate RNase L^36^.

Prior to Poly(I:C) transfection, the DFC exhibited its normal ring-like morphology. However, after Poly(I:C) transfection, FBL transformed into solid condensates in a population of cells (36.5%) (Figures 1G and 1H), indicating a disruption in the hollow shell structure of the DFC. In contrast, in cells expressing either BFP alone or the catalytically inactive RNase L R667A mutant, the ring-like morphology of FBL remained unchanged after Poly(I:C) transfection (Figures 1G and 1H). Our data indicate that RNA is essential for the formation of the hollow shell architecture of DFC in cells.

### rRNAs have specific localizations in the nucleolus

There are numerous RNAs in the nucleolus, including rRNAs. Mammalian cells contain four types of rRNAs: 5S, 5.8S, 18S, and 28S. The 5S rRNA is synthesized by RNA Polymerase III (Pol III) in the nucleoplasm, while the 5.8S, 18S, and 28S rRNAs are transcribed in the nucleolus by RNA Polymerase I (Pol I)^2^. Given that rRNA constitutes approximately 80% of the total RNA in cells, we hypothesized that rRNA is crucial for the formation of the DFC structure.

To explore this hypothesis, we investigated the localization of rRNAs in relation to the three sub-nucleolar compartments. We used the mNeonGreen-FBL stable cells to mark DFC and to delineate the locations of the GC and FC: the GC is located outside the FBL ring, and the FC is inside the FBL ring. We then conducted fluorescence in situ hybridization (FISH) in this cell line, using probes targeted to the 5’ETS of the 45S nascent pre-rRNA, and the mature forms of 5.8S, 18S, 28S, and 5S rRNA^38^ (Figure 2A).

**Figure 2.**
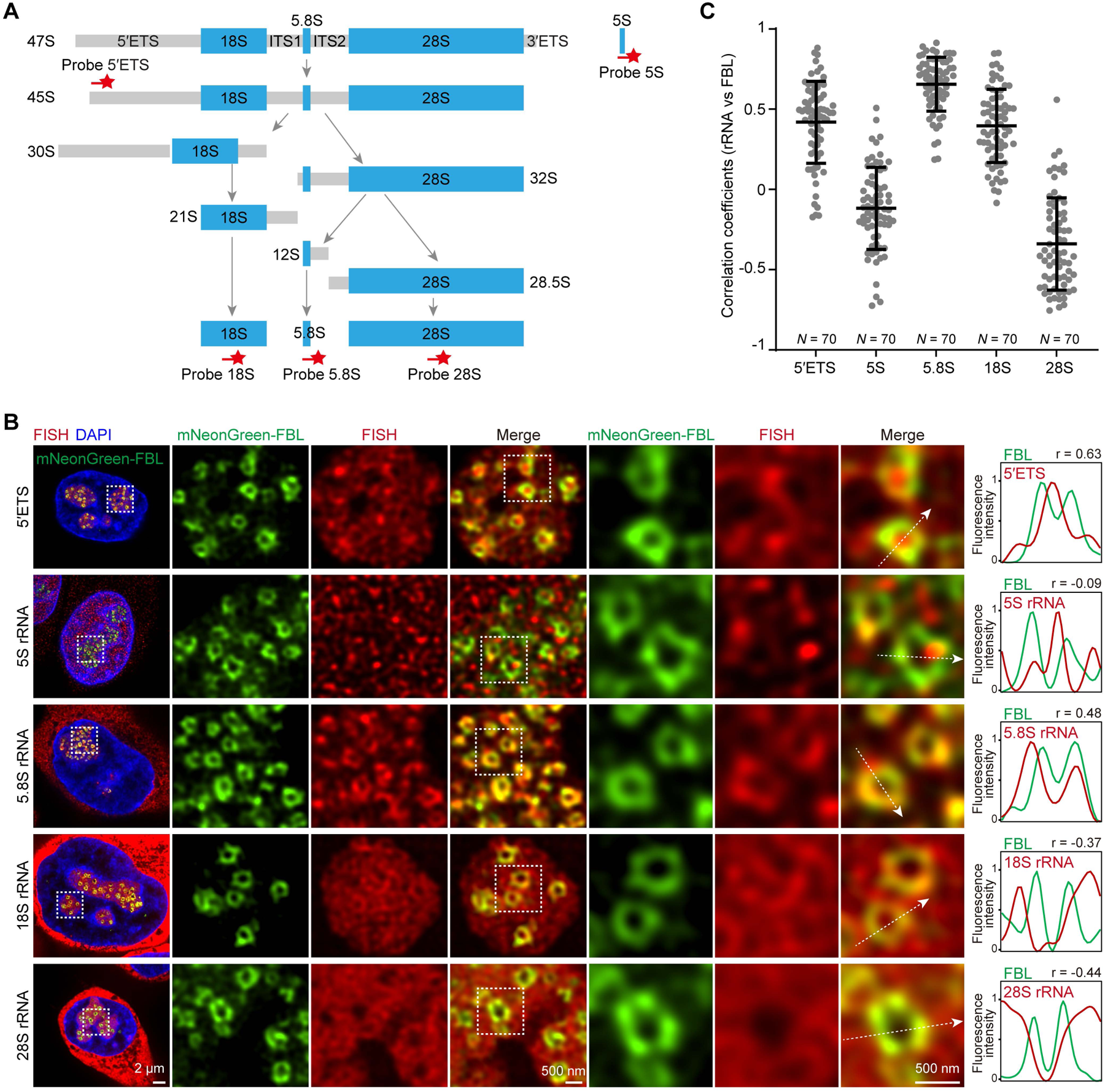
rRNAs have specific localizations in the nucleolus. (**A**) Schematic of a simplified pathway of rRNA processing and the location of probes used in the RNA-FISH experiment. (**B**) Representative super-resolution images of RNA-FISH targeting the 5’ETS, 5S, 5.8S, 18S, and 28S rRNA (red) in HeLa cells using probes in (A). mNeonGreen-FBL was stably expressed to label the nucleolar DFC. The nucleus was stained with DAPI. Shown are three different magnifications. Right: line profile of fluorescence intensities including Pearson’s correlation coefficients (r). The arrow indicates the plane used for the line profile generation. (**C**) Pearson’s correlation coefficients of line profiles of mNeonGreen-FBL and the indicated rRNAs from images in (B). The horizontal line denotes the median and error bars denote the 25th and 75th percentiles. *N*, number of all line profiles from three independent experiments.

Super-resolution microscopy revealed distinctive localization patterns for these rRNAs. The 45S pre-rRNA was localized exclusively in the nucleolus, with bright dots located within FBL rings (Figures 2B and S5A). This observation aligns with previous findings that pre-rRNA is transcribed at the FC-DFC interface^16^. In contrast, the 5S rRNA, although present in both the cytoplasm and nucleus, was not notably concentrated in the nucleolus (Figures 2B and S5A). The distribution of 5S rRNA appeared to be random and showed no specific enrichment in the DFC, as confirmed by line profile analysis and Pearson’s correlation coefficients (Figures 2B and 2C). The 5.8S, 18S, and 28S showed cytoplasm localizations and were strongly enriched in the nucleolus (Figures 2C and S5A), in agreement with that they are synthesized in the nucleolus.

Within the FC (inside the FBL ring), all three rRNAs exhibited decreased fluorescence signals (Figures 2B and 2C), consistent with the notion that rRNAs, once synthesized, are transferred to the DFC and GC for further processing. Notably, 5.8S, 18S, and 28S rRNA exhibited distinct localization patterns in relation to the DFC (Figures 2B and 2C). The 5.8S rRNA showed a high level of co-localization with the FBL ring (Figures 2B and 2C), indicating its enrichment in the DFC (Figures 2B and 2C). The 18S rRNA showed partial co-localization with the FBL ring (Figures 2B and 2C) but was not as enriched as that of 5.8S. The 28S rRNA was highly concentrated in the GC (outside the FBL ring) and partially co-localized with the FBL ring, with a decreased signal towards the FC. This pattern led to the formation of assemblies with hole-like structures encircling the DFC (Figures 2B and 2C). Since 5.8S, 18S, and 28S rRNA are generated from the cleavage of a single pre-rRNA, the mechanisms contributing to their differential localization remain unknown.

### In vitro reconstitution of DFC-like structures with 28S rRNA

Having observed the close connection between rRNA localizations and the DFC, we tested their roles in the DFC structure formation. By treating cells with CX-5461, a Pol I inhibitor, we aimed to inhibit rRNA production in the nucleolus. We observed a significant disruption in the ring-like structure of FBL (Figure 3A), suggesting that rRNAs are essential for the formation of hollow shell structures of DFC in cells.

**Figure 3.**
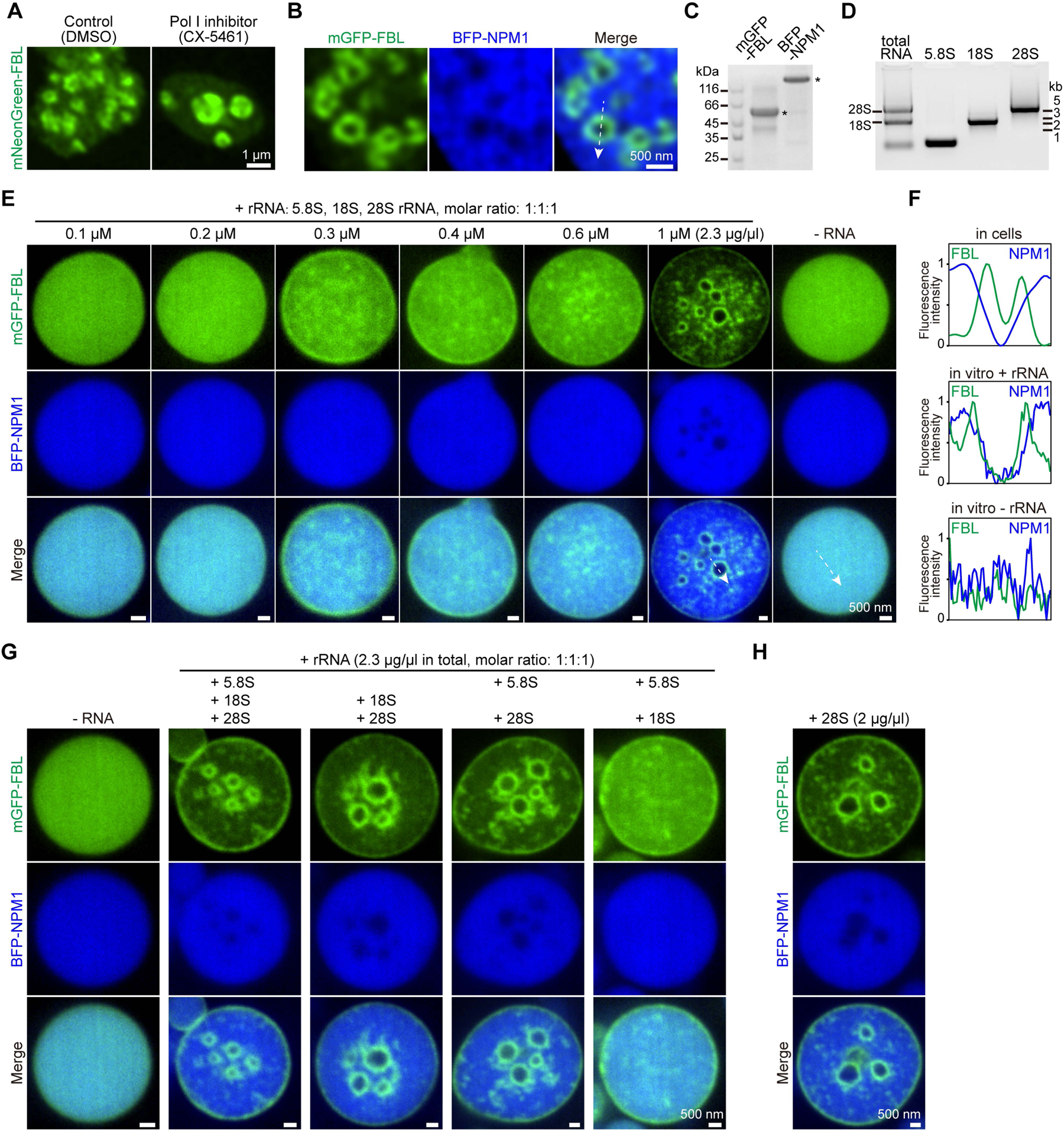
In vitro reconstitution of DFC-like structures with 28S rRNA. (**A**) Representative live-cell imaging of nucleoli in HeLa cells treated with DMSO or CX-5461(1 µM). mNeonGreen-FBL was stably expressed to label the nucleolar DFC. (**B**) Representative live-cell imaging of nucleoli in HeLa cells. mGFP-FBL and BFP-NPM1 were transiently transfected into cells to label nucleolar DFC and GC respectively. The line profile of fluorescence intensities of FBL and NPM1 was shown in (F). The arrow indicates the plane used for the line profile generation. (**C**) SDS-PAGE of the purified mGFP-tagged FBL and BFP-tagged NPM1 protein. The star indicates the targeted band. (**D**) Agarose gel electrophoresis of the total RNA extracted from HeLa cells and the in vitro transcribed full-length human 5.8S, 18S, 28S rRNA. In vitro transcribed rRNAs exhibited the same molecular weight as the rRNAs extracted from HeLa cells. (**E**) Representative confocal images of phase separation experiments using purified mGFP-FBL (5 µM) and BFP-NPM1 (30 µM) in the absence or presence of human 5.8S, 18S, and 28S rRNA with the indicated concentrations (1 µM means 1 µM 5.8S + 1 µM 18S + 1 µM 28S, the concentration of all three rRNAs is 2.3 µg/µl). The line profile of fluorescence intensities of FBL and NPM1 was shown in (F). The arrow indicates the plane used for the line profile generation. (**F**) line profile of fluorescence intensities of FBL and NPM1 from (B) (in cells) and (E) (in vitro). (**G**) same as (E), but the concentration of total rRNAs is kept constant (2.3 µg/µl). (**H**) same as (E).

In human cells, hundreds of rRNA gene copies are located in clusters of tandem repeats across five chromosomes^2^. Knocking out or knocking down the expression of specific rRNAs in cells is technically challenging. To carefully assess the role of rRNA in DFC formation, we employed an established in vitro nucleolar reconstitution approach^28^. We first purified mGFP-tagged FBL and BFP-tagged NPM1, which are scaffold proteins for the DFC and GC, respectively (Figures 3B and 3C). We then cloned human 5.8S, 18S, and 28S genes and synthesized full-length rRNAs using the T7 in vitro transcription system (Figure 3D). Notably, the 5S rRNA was not included in our reconstitution experiments, as it is not enriched in the nucleolus (Figure 2B). We performed phase separation experiments by mixing mGFP-FBL and BFP-NPM1, either without or with the addition of rRNAs. We noted that in the absence of rRNA, FBL and NPM1 co-phase separated to form condensates displaying uniformly distributed FBL or NPM1 proteins (Figures 3E, 3F, and S6A). Strikingly, upon adding 5.8S, 18S, and 28S rRNAs, and upon reaching a critical concentration, we observed the formation of multiple ring-like structures of FBL within the condensate (Figures 3E, 3F, and S6A). These structures largely excluded NPM1 internally (Figures 3E and 3F), which is similar to the localization pattern of FBL and NPM1 in cells, indicating formation of hollow shell DFC-like structures (Figures 3B and 3F). These observations, along with the results of the Pol I inhibition experiment (Figure 3A), confirm that rRNAs are critical for inducing the hollow shell structure of DFC.

Next, we sought to identify which specific rRNA was necessary for this process. We conducted in vitro reconstitution with various combinations of two out of the three rRNAs. Remarkably, the omission of 28S, rather than 5.8S or 18S rRNA, prevented the formation of FBL hollow shells (Figure 3G). Moreover, adding 28S rRNA alone, but not 5.8S or 18S, successfully induced the formation of FBL hollow shells in a concentration-dependent manner (Figures 3H and S7A–S7C). Therefore, we concluded that 28S rRNA is the key component responsible for the formation of the hollow shell structure of DFC.

### Multivalent interactions of 28S rRNA are required to induce the formation of DFC-like structures

To elucidate the mechanism behind the formation of DFC-like structures, we first examined whether adding 28S rRNA to FBL or NPM1 would be sufficient to induce hollow shell structures. When mixed without or with 28S rRNA, FBL formed small condensates exhibiting a beads-on-a-string arrangement (Figure 4A). On its own, NPM1 formed spherical condensates. The addition of 28S rRNA enhanced NPM1 phase separation, as evidenced by the increased size of the NPM1 condensates (Figure 4A). Notably, the formation of FBL hollow shell structures, with NPM1 exclusion inside the shell, was only observed when FBL, NPM1, and 28S rRNA were combined (Figure 4A). These observations indicate that interactions among the GC component, the DFC component, and the 28S rRNA is essential for the formation of DFC-like structures.

**Figure 4.**
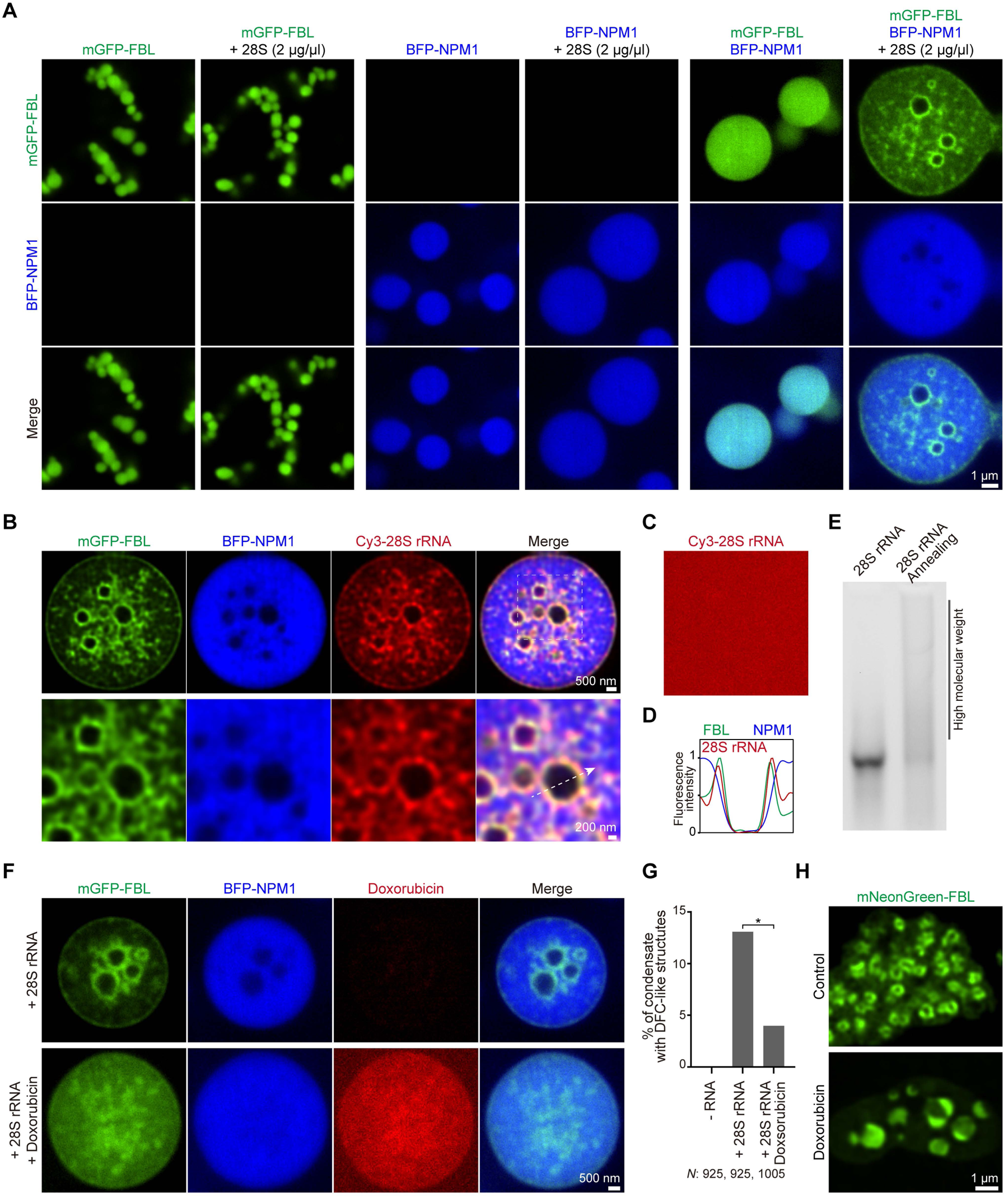
Multivalent interactions of 28S rRNA are required to induce the formation of DFC-like structures. (**A**) Representative confocal images of phase separation experiments using purified mGFP-FBL (5 µM) or BFP-NPM1 (30 µM) in the absence or presence of 28S rRNA (2 µg/µl). (**B**) the same as (A), but with Cy3 labeled 28S rRNA. The line profile of fluorescence intensities of FBL, NPM1, and 28S rRNA was shown in (D). The arrow indicates the plane used for the line profile generation. (**C**) Representative confocal images of Cy3 labeled 28S rRNA (2 µg/µl) in the phase separation buffer. (**D**) The line profile of fluorescence intensities from (B). (**E**) Native agarose gel electrophoresis of the in vitro transcribed 28S rRNA before (lane 1) and after RNA interaction experiment (Annealing) (lane 2). (**F**) the same as (A), but 28S rRNA was pretreated without or with Doxorubicin (250 µM) for two hours. (**G**) Quantification of FBL-NPM1 condensates with the formation of hollow shell DFC-like structures in (F). *N*, the total number of condensates in each condition from three independent experiments. t test, * p < 0.05. (**H**) Representative live-cell imaging of nucleoli in HeLa cells treated without or with Doxorubicin (2.5 µM). mNeonGreen-FBL was stably expressed to label the nucleolar DFC.

Next, we synthesized Cy3-labeled 28S rRNA to visualize its localization (Figures 4B and 4C). In the phase separation buffer, 28S rRNA alone displayed a uniform fluorescence signal (Figure 4C), indicating that 28S rRNA did not form any detectable assemblies by itself. In contrast, within the FBL-NPM1 condensates, 28S rRNA displayed an uneven fluorescence signal distribution (Figures 4B and 4D). 28S rRNA primarily localized with NPM1 and was enriched at the FBL ring, but it was excluded from the inside of the FBL ring (Figures 4B and 4D). This pattern resembles the localization of 28S rRNA in the nucleolus: excluded from the FC, co-localized with the DFC shell, and extensively present in the GC (Figures 2B and 2C). These data demonstrate that 28S rRNA forms assemblies within the condensates.

Given that multivalent RNAs have been reported to form intermolecular interactions to assemble as the skeleton of RNA granules^35^, we hypothesized that multivalent interactions of 28S rRNA are crucial for their ability to induce DFC-like structures. To test this, we conducted RNA interaction experiments. The detection of high molecular weight 28S rRNA in native agarose gel electrophoresis confirmed extensive RNA-RNA interactions (Figure 4E), verifying the multivalency of 28S rRNA. Next, we treated 28S rRNA with doxorubicin, a nucleic acid intercalator, before setting up the phase separation experiment to perturb RNA-RNA interactions. Doxorubicin treatment effectively prevented the formation of 28S rRNA-induced DFC-like structures in vitro (Figures 4F and 4G). Moreover, doxorubicin treatment potently disrupted the hollow shell structures of DFC in cells (Figure 4H). In summary, our results indicate that the multivalent interactions of 28S rRNA are necessary for the induction of DFC’s hollow shell structures.

### 28S rRNAs from organisms with a tripartite nucleolus induce the formation of hollow shell DFC-like structures

Building on our findings, we developed a hypothesis that 28S rRNA plays a key role in the transition from a bipartite to a tripartite nucleolar structure. If this hypothesis holds true, then 28S rRNA from other organisms with a tripartite nucleolus should also be capable of inducing hollow shell (DFC-like) structures in the in vitro nucleolar reconstitution experiment. To test this theory, we cloned the 28S rRNA genes from three model organisms known to have tripartite nucleoli: mouse, chicken^9^, and baker’s yeast^12,13^ (Saccharomyces cerevisiae) (which has 25S rRNA) (Figure 5A). As a control, we also cloned the 28S rRNA genes from two model organisms with bipartite nucleoli: Drosophila^14^ and C. elegans^15^ (which possess 26S rRNA) (Figure 5A). We then synthesized these 28S rRNAs (Figure 5B) and conducted in vitro nucleolar reconstitution experiments.

**Figure 5.**
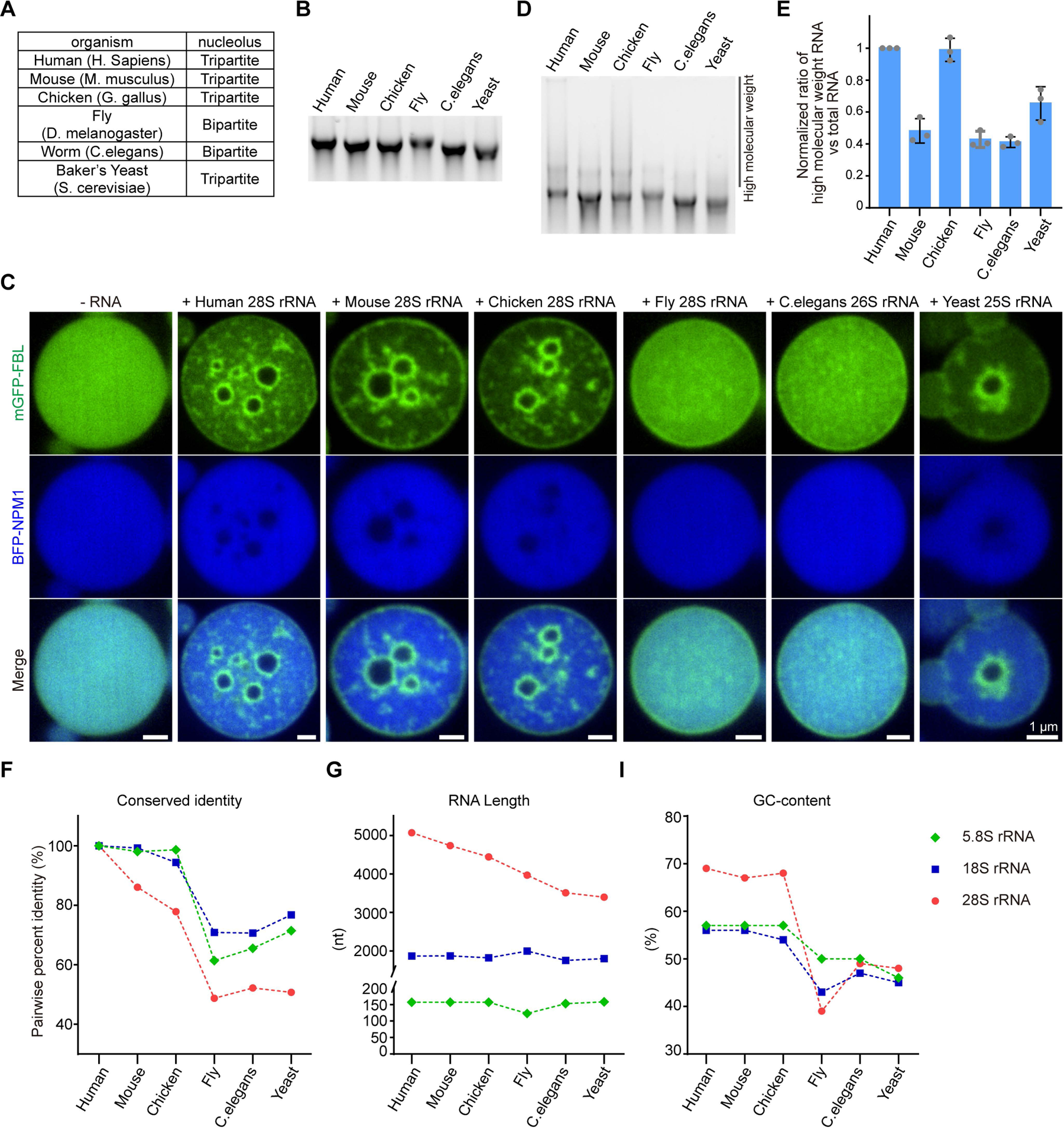
28S rRNAs from organisms with a tripartite nucleolus induce the formation of hollow shell DFC-like structures. (**A**) Schematic showing the type of nucleoli of each model organism. (**B**) Agarose gel electrophoresis of the in vitro transcribed 28S rRNAs of the indicated organisms. (**C**) Representative confocal images of phase separation experiments using purified mGFP-FBL (5 µM) and BFP-NPM1 (30 µM) in the absence or presence of 28S rRNA (2 µg/µl) of the indicated organisms. (**D**) Native agarose gel electrophoresis of the in vitro transcribed 28S rRNA of the indicated organisms after RNA interaction experiment. (**E**) Quantification of the extent of high molecular weight RNA species formed by 28S rRNAs of the indicated organisms in (D). human 28S rRNA was normalized to 1. Shown is mean ± SD of three independent experiments. (**F**) Pairwise percent identity of the 28S rRNA of the indicated organisms vs human 28S rRNA. (**G**) Length of 28S rRNAs of the indicated organisms. (**H**) Pairwise sequence alignment of human 28S and C.elegans 26S rRNA. Shown is a fragment of extended sequences in the human 28S flanked by conserved sequences. Stars indicate identical nucleotides. G and C are marked red within the extended sequences. (**I**) GC-content of 28S rRNAs of the indicated organisms.

Consistent with our hypothesis, the 28S rRNAs from all the tripartite nucleolus organisms tested successfully induced the formation of hollow shell DFC-like structures (Figure 5C). In contrast, 28S rRNAs from the bipartite nucleolus organisms failed to induce the formation of DFC-like structures (Figure 5C). These observations support our hypothesis that 28S rRNA plays a crucial role in the transition of the nucleolar structure from bipartite to tripartite.

### Emerging of multivalency of 28S rRNAs during evolution

Given that multivalent interactions of human 28S rRNA are essential for inducing hollow shell DFC-like structures, we wondered whether there are variations in the degree of multivalency among 28S rRNAs. To explore this, we carried out RNA interaction experiments along with native agarose gel electrophoresis (Figure 5D). These experiments revealed significant differences among 28S rRNAs in the formation of high molecular weight RNA complexes, indicating varying extents of multivalent RNA-RNA interactions (Figures 5D and 5E). Notably, 28S rRNAs from organisms with a tripartite nucleolus (human, chicken, mouse, and baker’s yeast) exhibited higher levels of multivalent interactions (Figures 5D and 5E). In contrast, 28S rRNAs from bipartite nucleolus organisms showed the lowest levels of multivalency (Figures 5D and 5E). In summary, our data suggest that during evolution, 28S rRNA in tripartite nucleolus organisms has acquired a greater degree of multivalency, which correlates with its ability to induce the formation of hollow shell DFC-like structures.

Next, we moved to identify features of 28S rRNA that contribute to its multivalency. We first aligned the 5.8S, 18S, and 28S rRNA sequences from six model organisms to compare their conservation score (Table S1 to S3). Strikingly, 28S rRNA showed lower conservation across species compared to 5.8S and 18S rRNAs (Figure 5F). For example, the pairwise percent identity was 99.3% for 18S but only 86.1% for 28S between mouse and human, and 70.7% for 18S but 52.2% for 28S between C. elegans and human (Figure 5F). Additionally, 28S rRNAs in higher organisms have substantially increased in length, while the lengths of 5.8S and 18S rRNAs remain relatively constant (Figure 5G). Furthermore, the extended sequences in 28S rRNA of higher organisms tend to be GC-rich (Figures 5H, S8A, S8B, and Table S3), leading to a higher overall GC-content compared to that in lower organisms (Figure 5I). Interestingly, the increased GC-content is also observed in 5.8S and 18S rRNAs (Figure 5I). It has been reported that longer RNAs have a greater capacity for forming intermolecular RNA-RNA interactions^39^. In addition, GC-rich RNAs can form more complex interaction types, such as G-quadruplexes, besides canonical base-pairing^30^. Thus, the increased length and higher GC-content are key factors contributing to the enhanced multivalency of 28S rRNA. Interestingly, despite the 25S rRNA of baker’s yeast having a similar length and GC-content to the C. elegans 26S rRNA, it exhibited a higher degree of multivalent RNA-RNA interactions (Figures 5D and 5E). Similarly, despite mouse 28S and chicken 28S rRNA having comparable lengths and GC contents, the mouse 28S rRNA displayed a lower level of multivalency (Figures 5D and 5E). These observations indicate that additional factors beyond length and GC content contribute to the multivalency of 28S rRNA.

### 28S rRNA induces the formation of three-layered nucleolar GC-DFC-FC-like structures

We posited that 28S rRNA-induced formation of the hollow shell structure of DFC is a critical step in generating a three-layered nucleolus, as it acts as a barrier separating the GC and the FC, which is located inside the DFC shell. To experimentally investigate this hypothesis, we examined whether 28S rRNA can induce the formation of three-layered GC-DFC-FC-like structures in the reconstitution experiment. For this purpose, we purified mCherry-tagged POLR1D (Figure 6A), a subunit of RNA Polymerase I localized in the nucleolar FC (Figure 6B)^17^, and then performed in vitro nucleolar reconstitution. In experiments mixing NPM1, FBL, and POLR1D with human 28S rRNA, we observed the formation of three-layered structures within the condensate, characterized by enriched POLR1D localization and NPM1 exclusion inside the FBL shell (Figure 6C). This arrangement closely resembles the nested GC-DFC-FC layers observed in the tripartite nucleolus (Figure 6B). Furthermore, 28S rRNAs from other organisms with a tripartite nucleolus also successfully induced similar three-layered GC-DFC-FC-like structures (Figure 6C). In contrast, 28S rRNAs from organisms with a bipartite nucleolus did not lead to the formation of these three-layered structures (Figure 6C). In summary, we conclude that multivalent 28S rRNA is instrumental in inducing the formation of three-layered nucleolar structures resembling GC-DFC-FC.

**Figure 6.**
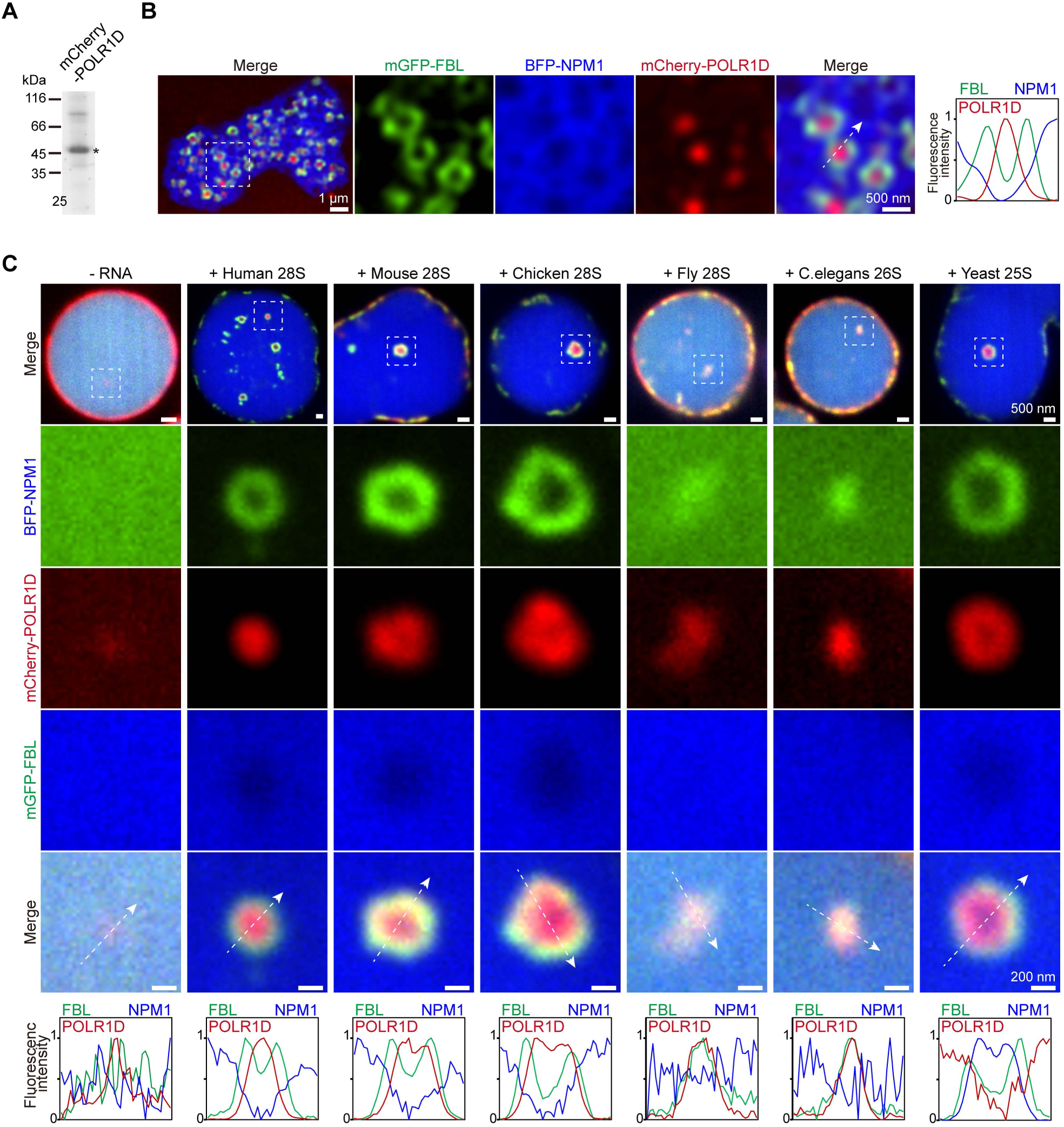
28S rRNA induces the formation of three-layered nucleolar GC-DFC-FC-like structures. (**A**) SDS-PAGE of the purified mCherry-tagged POLR1D protein. The star indicates the targeted band. (**B**) Representative live-cell imaging of nucleoli in HeLa cells. mGFP-FBL, BFP-NPM1, and mCherry-POLR1D were transiently transfected into cells to label nucleolar DFC, GC, and FC, respectively. Shown are two different magnifications. Right: line profile of fluorescence intensities. The arrow indicates the plane used for the line profile generation. (**C**) Representative confocal images of phase separation experiments using purified mGFP-FBL (5 µM), BFP-NPM1 (30 µM), and mCherry-POLR1D (30 µM), in the absence or presence of 28S rRNA (2 µg/µl) of the indicated organisms. Bottom: line profiles of fluorescence intensities. The arrow indicates the plane used for the line profile generation.

## Discussion

rRNA constitutes 80% of the total RNA in a single cell. The 28S rRNA is the longest, with a molecular weight of 1650 kd in humans, significantly larger than 18S (609 kd), 5.8S (51 kd), and 5S (40 kd) rRNAs. Thus, by weight, 28S accounts for about 70% of rRNA and 56% of the total RNA, making it the most abundant RNA in cells. While rRNA genes are among the most conserved genes^40^, 28S rRNA, despite being closely located with 18S and 5.8S in the same transcription units, is less conserved than 18S and 5.8S. In higher organisms, 28S has undergone substantial lengthening along with an increase in GC content, raising intriguing questions about how these additional properties were acquired. The distance between the 28S and 5.8S genes is relatively short, at 1167 bp, yet the factors driving the specific lengthening of 28S remain unknown. Furthermore, the eukaryotic genome contains hundreds of tandem 18S-5.8S-28S gene units clustered on one or several chromosomes^2^. How such dramatic changes in the sequence of 28S are spread and coordinated across all these units presents another intriguing question. Nevertheless, our research has demonstrated that the lengthening, higher GC content, and other yet-to-be-identified features endow 28S rRNA with greater multivalency in organisms with a tripartite nucleolus. Our in vitro nucleolar reconstitution experiments suggest that the emergence of 28S rRNA’s multivalency facilitated the transition from a bipartite to a tripartite nucleolus.

Despite these findings, it is important to note that the three-layered nucleolar-like structures formed in our reconstitution experiments do not exactly replicate the nucleolus. For example, the DFC-like structures are formed in only a portion of FBL-NPM1 condensates (Figures 4G and S6A), and the number of DFC-like structures formed in vitro (In a portion of FBL-NPM1 condensates, only one DFC-like structure was formed) is less than the number of DFC in the HeLa nucleolus; in the DFC-like structure with a large size, POLR1D localizes close to the rim of the FBL ring instead of filling the FBL shell (Figure S9A). One reason for these differences might be that the concentration of 28S rRNA used in our reconstitution is lower than that in the nucleolus. Limited by the yield of in vitro transcription and the solubility of 28S rRNA, we used up to about 2 µg/µl rRNAs in the reconstitution. In HeLa cells, the RNA concentration in the nucleus is reported to be around 8.5 µg/µl^41^. Considering most RNAs are rRNAs, and nuclear rRNAs are primarily localized in the nucleolus (Figures 2B and S5A), the actual rRNA concentration in the nucleolus could be one order of magnitude higher than that used in the reconstitution. Additionally, the nucleolus is composed of thousands of proteins and numerous RNA species, with many factors contributing to its complex architecture. For instance, TCOF1 is involved in FC formation^15^; *SLERT*, a long non-coding RNA, and DDX21, an RNA helicase, regulate DFC size^18^. More recently, a new layer, the PDFC, was identified between the DFC and GC in the nucleolus^17^. In addition, the nucleolus was found to exhibit much more complex material properties than the typical liquid^42^, and the micropolarity regulates the structure of the nucleolus^43^.

The in vitro nucleolar reconstitution provides a bottom-up platform for future exploration into new factors that regulate nucleolar structures and properties.

## Acknowledgments

We thank all members of the Ma lab for helpful discussions. We thank Christine Mayr (Memorial Sloan Kettering Cancer Center) for providing HeLa cell and pcDNA vectors. We thank Long Zhang (Zhejiang University) for providing pLVX vectors. We thank Tong Chao (Zhejiang University), Cunqi Ye (Zhejiang University), and Xin Li (Zhejiang University) for providing Fly RNA, Yeast RNA, and Chicken RNA, respectively. We thank Jingxiu Xu (Zhejiang University) and Suhong Xu (Zhejiang University) for providing C.elegans cDNA. We thank Jian Zhang (Yunnan University) and Jinghao Sheng (Zhejiang University) for critically reading the manuscript and suggestions.

This work was funded by the National Natural Science Foundation of China (32370727), the Leading innovation and entrepreneurship team of Hangzhou (TD2020006), the start-up funding from the Life Sciences Institute to W.M. and the National Key R&D Program (2019YFA0508403), the National Natural Science Foundation of China (32090044, and 31971128), the Strategic Priority Research Program of the Chinese Academy of Sciences (XDB 37040202), the CAS Project for Young Scientists in Basic Research (YSBR-068) to S.X..

## Author Contributions

W.M. JW and S.X. conceived the projects and designed the experiments. J.W. performed most of phase separation experiments, experiments with mammalian cells and analyses. Y.Z. purified recombinant proteins. X.F. cloned the yeast 25S rRNA. Y.H. cloned the chicken 28S rRNA and the fly 26S rRNA. B.Y. performed the western blot experiment and provided help with line profile analysis. W.M. and J.W. wrote the manuscript.

## Supplementary Figures

**Figure S1.**
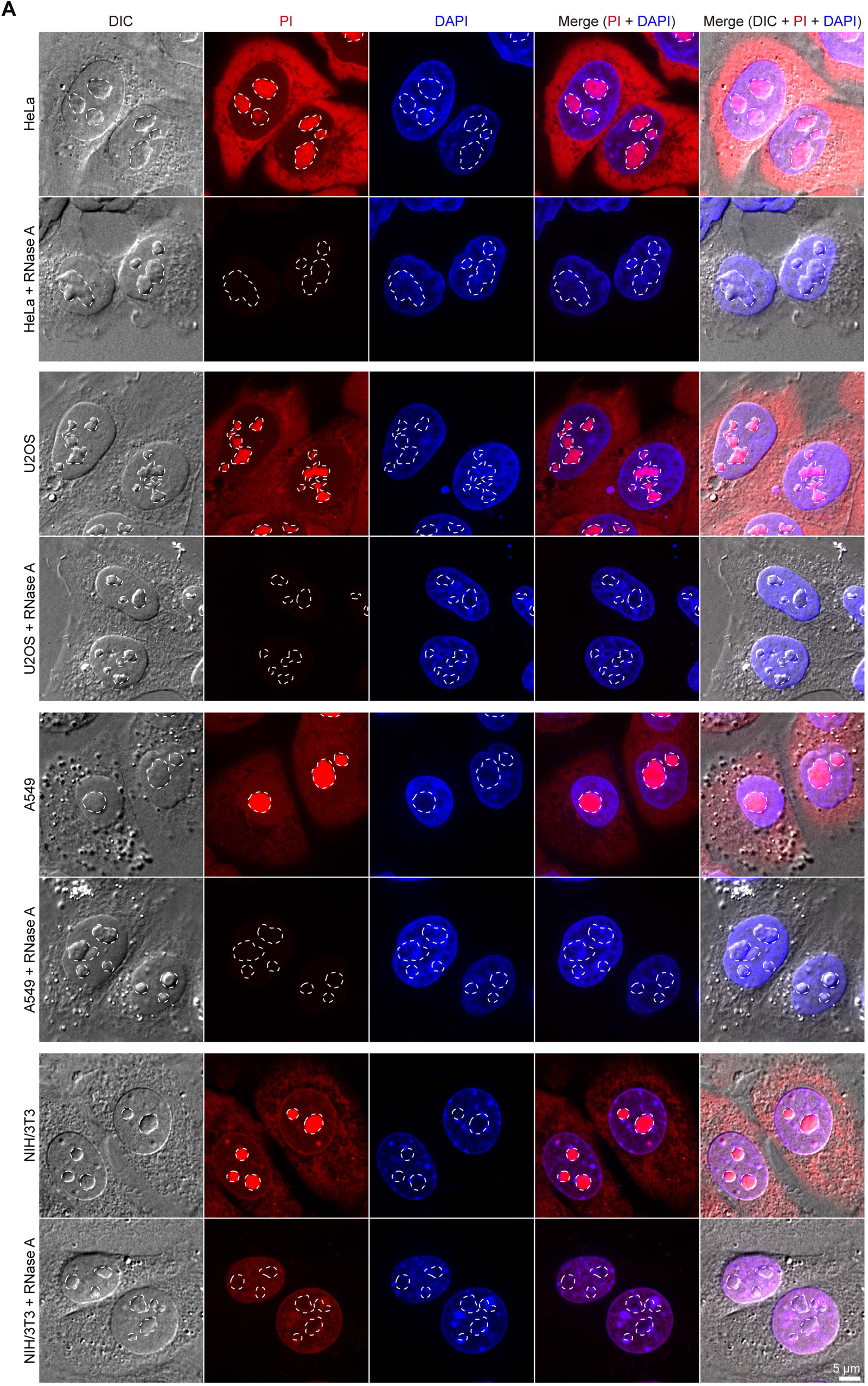
The nucleolus contains a high concentration of RNA. (**A**) Representative confocal images of HeLa, U2OS, A549, and NIH/3T3 cells stained with PI and DAPI. To degrade RNA, cells were treated with RNase A before the PI staining. The nucleolus is indicated by DIC imaging and is marked by the white dotted line.

**Figure S2.**
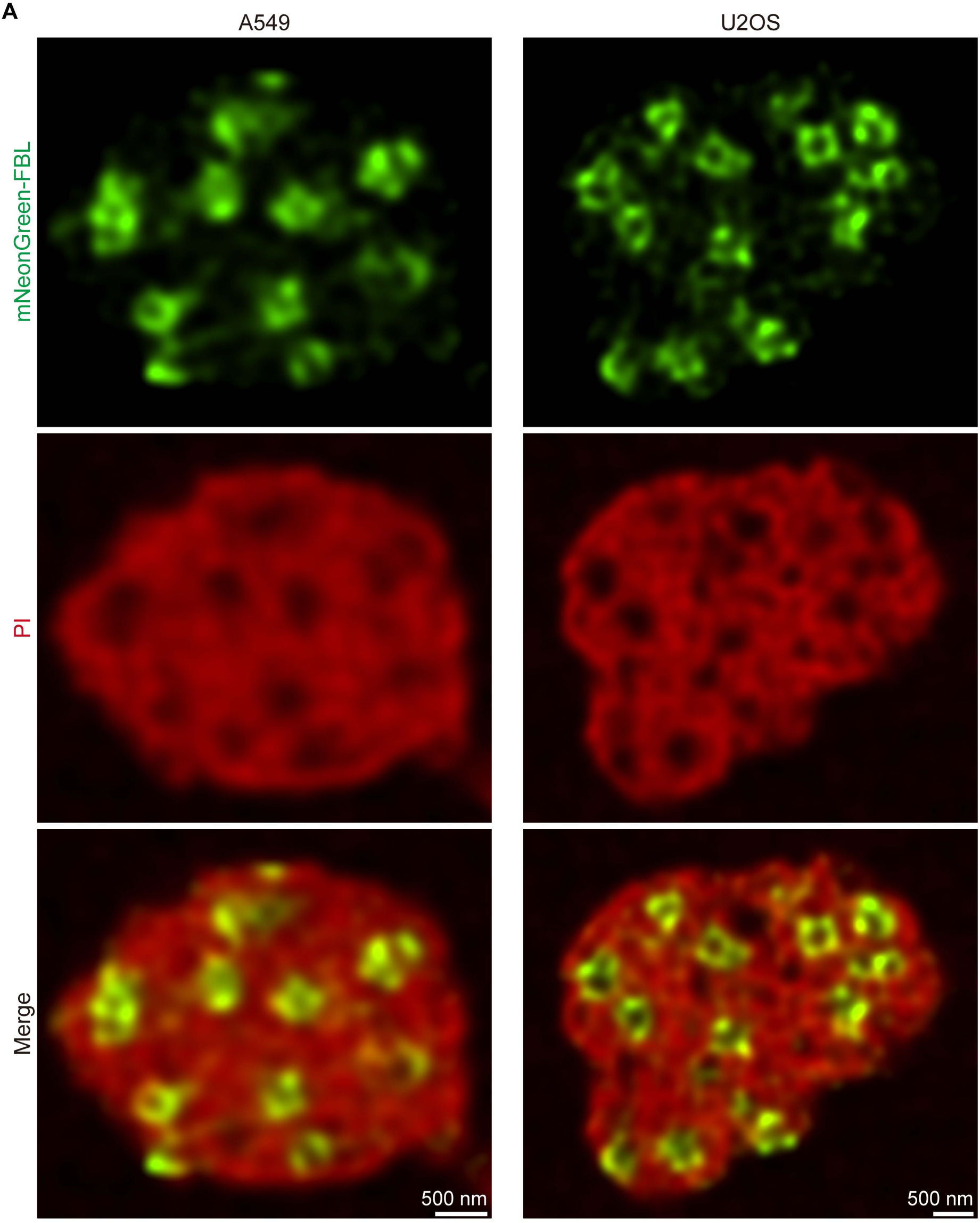
RNA forms assemblies surrounding the DFC in the nucleolus. (**A**) Representative super-resolution images of the nucleolar DFC and RNA (PI staining) in U2OS and A549 cells. Stably expressed mNeonGreen-FBL labels the DFC.

**Figure S3.**
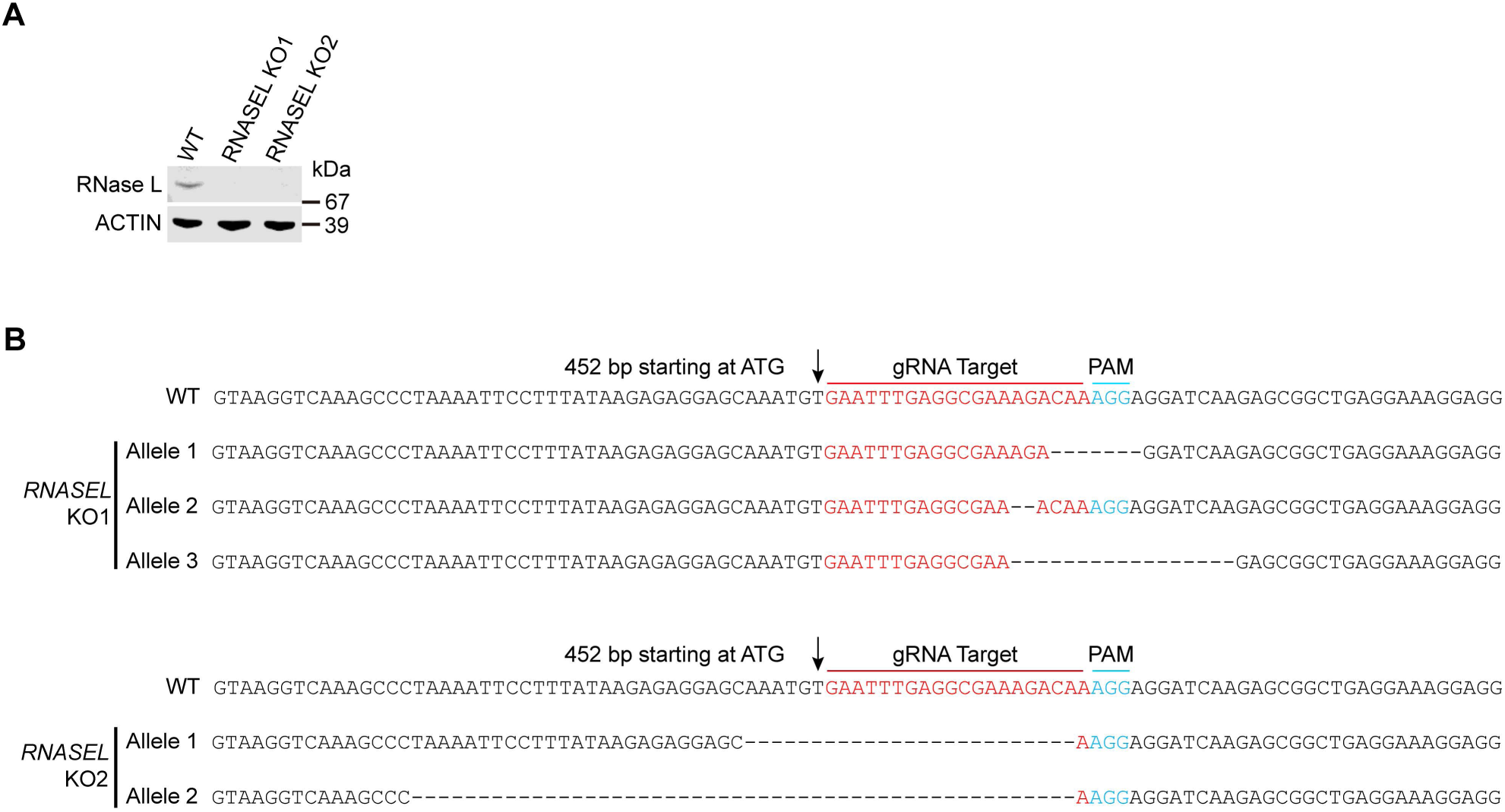
Generation of *RNASEL* knockout cells. (**A**) Western blot of wild-type (WT), and *RNASEL* knockout HeLa cells using the indicated antibodies. (**B**) *RNASEL* genomic locus showing the wild-type and the frame-shift mutations introduced using the CRISPR/Cas9 system with the indicated guide RNA.

**Figure S4.**
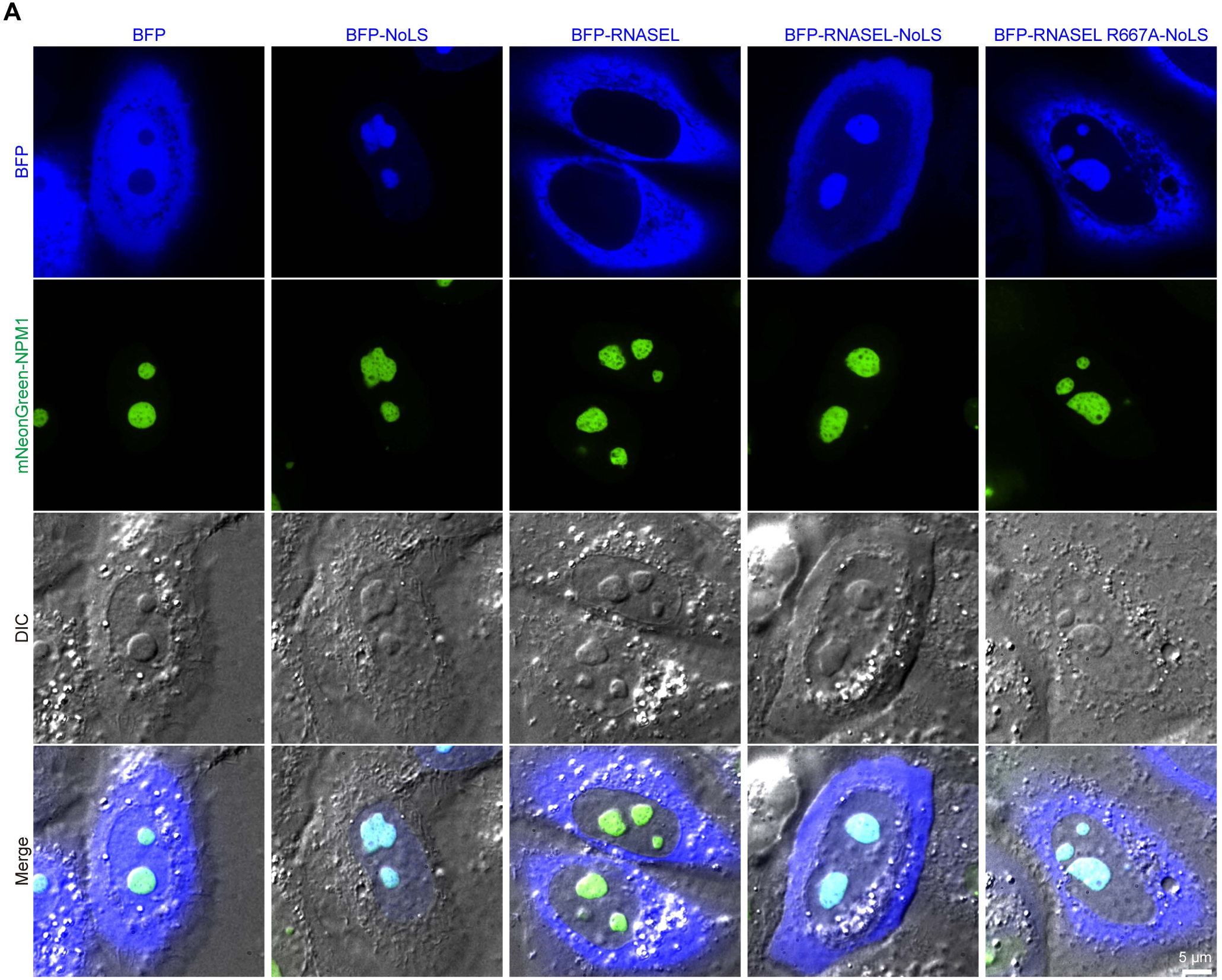
The NoLS targets RNase L to the nucleolus. (**A**) Representative live cells images of HeLa cells transiently transfected with the indicated fusion proteins. mNeonGreen-NPM1 labels the nucleolus. 3x tandem repeats of SV40 NLS (PKKKRKV) (nuclear localization signal) function as a NoLS (nucleolus localization signal) to target RNase L to the nucleolus.

**Figure S5.**
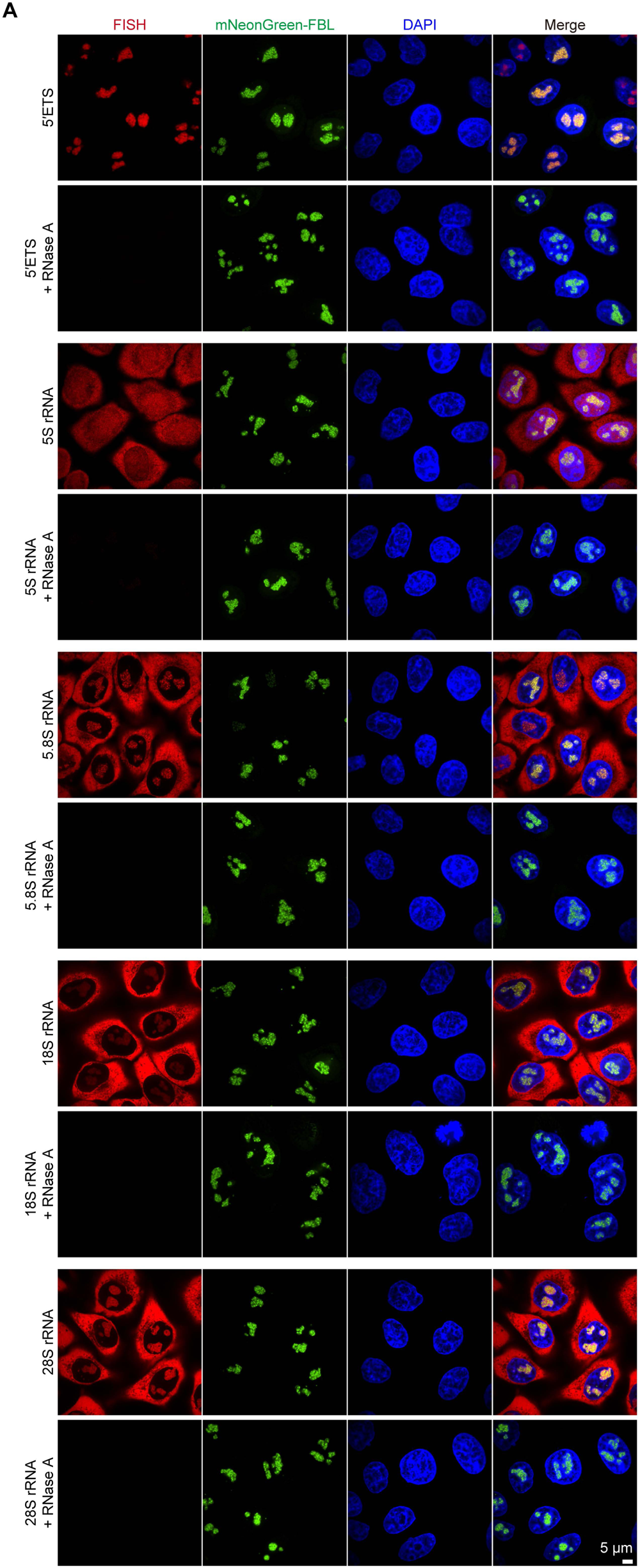
Fluorescence in situ hybridization (FISH) of rRNAs. (**A**) Representative confocal images of RNA-FISH targeting the 5’ETS, 5S, 5.8S, 18S, and 28S rRNA (red) in HeLa cells using probes in Fig. 2A. To degrade RNA, cells were treated with RNase A. Stably expressed mNeonGreen-FBL labels the DFC. The nucleus was stained with DAPI.

**Figure S6.**
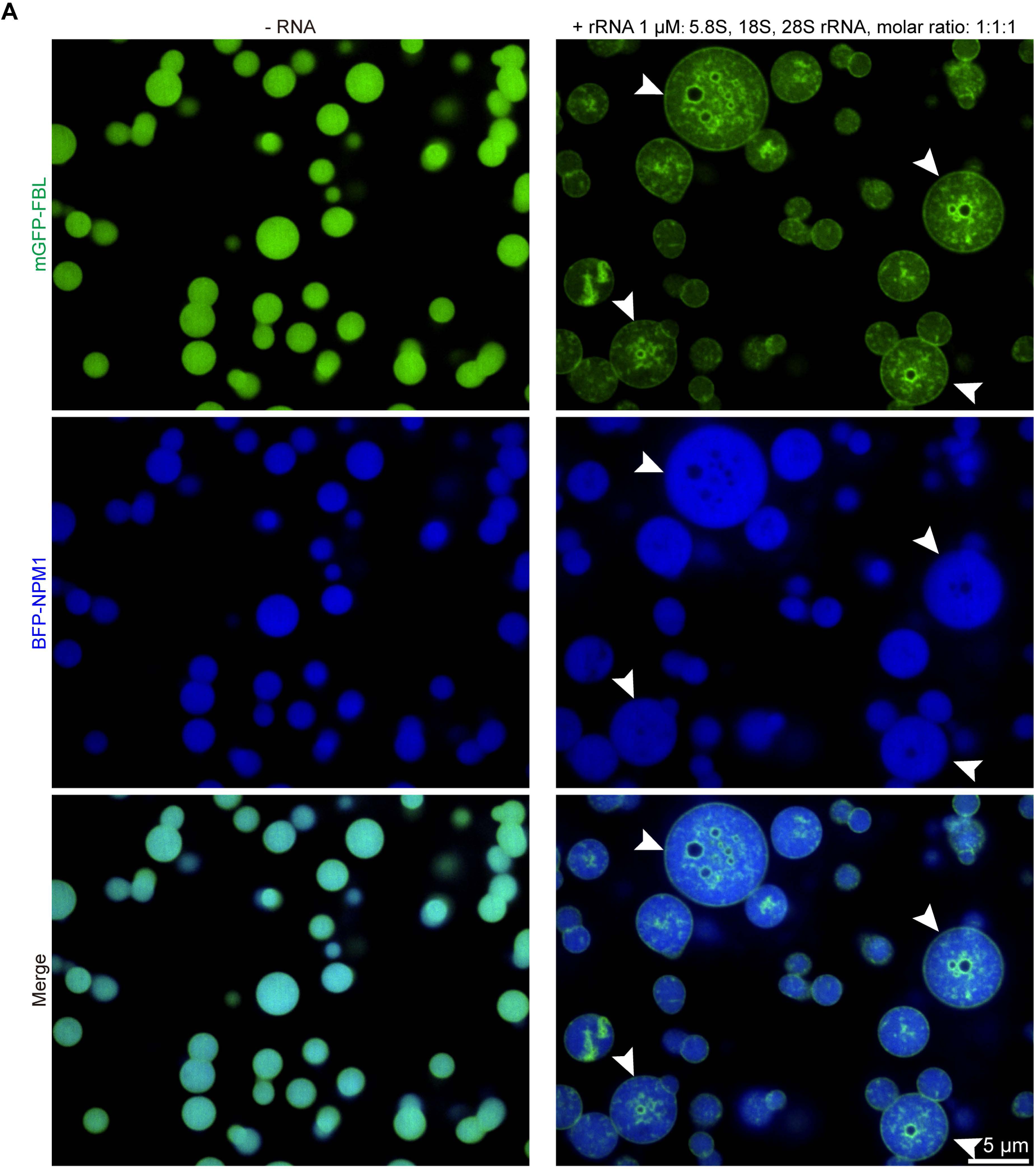
rRNAs induce the formation of DFC-like structures in vitro. (**A**) Representative confocal images of phase separation experiments using purified mGFP-FBL (5 µM) and BFP-NPM1 (30 µM) in the absence or presence of the indicated in vitro transcribed rRNAs. Arrowheads indicate FBL-NPM1 with hollow shell DFC-like structures.

**Figure S7.**
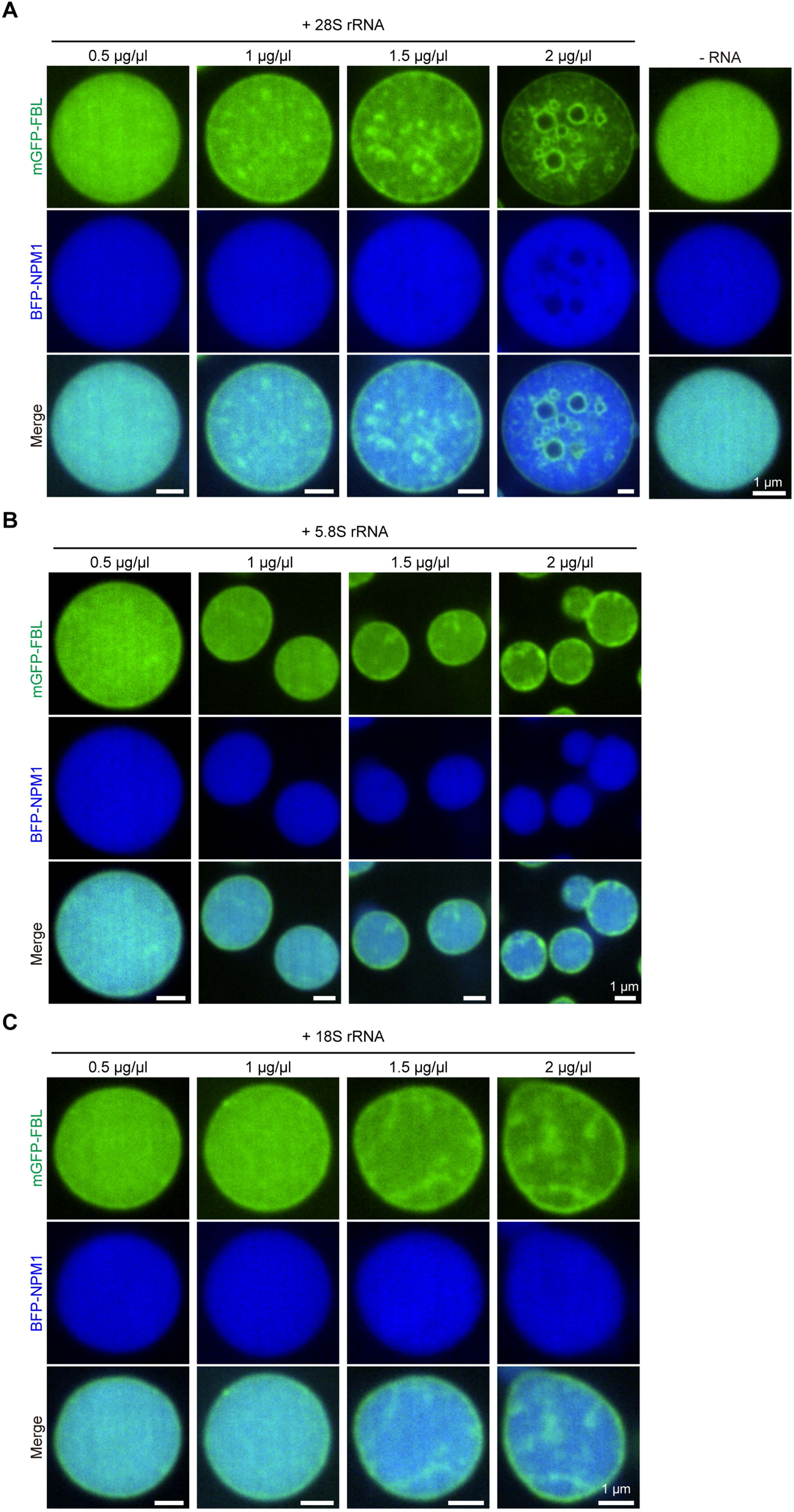
In vitro reconstitution of DFC-like structures with 28S rRNA. (**A**) Representative confocal images of phase separation experiments using purified mGFP-FBL (5 µM) and BFP-NPM1 (30 µM) in the absence or presence of the in vitro transcribed 28S rRNA with the indicated concentration. (**B**) as in (A), but in the presence of 5.8S rRNA. (**C**) as in (A), but in the presence of 18S rRNA.

**Figure S8.**
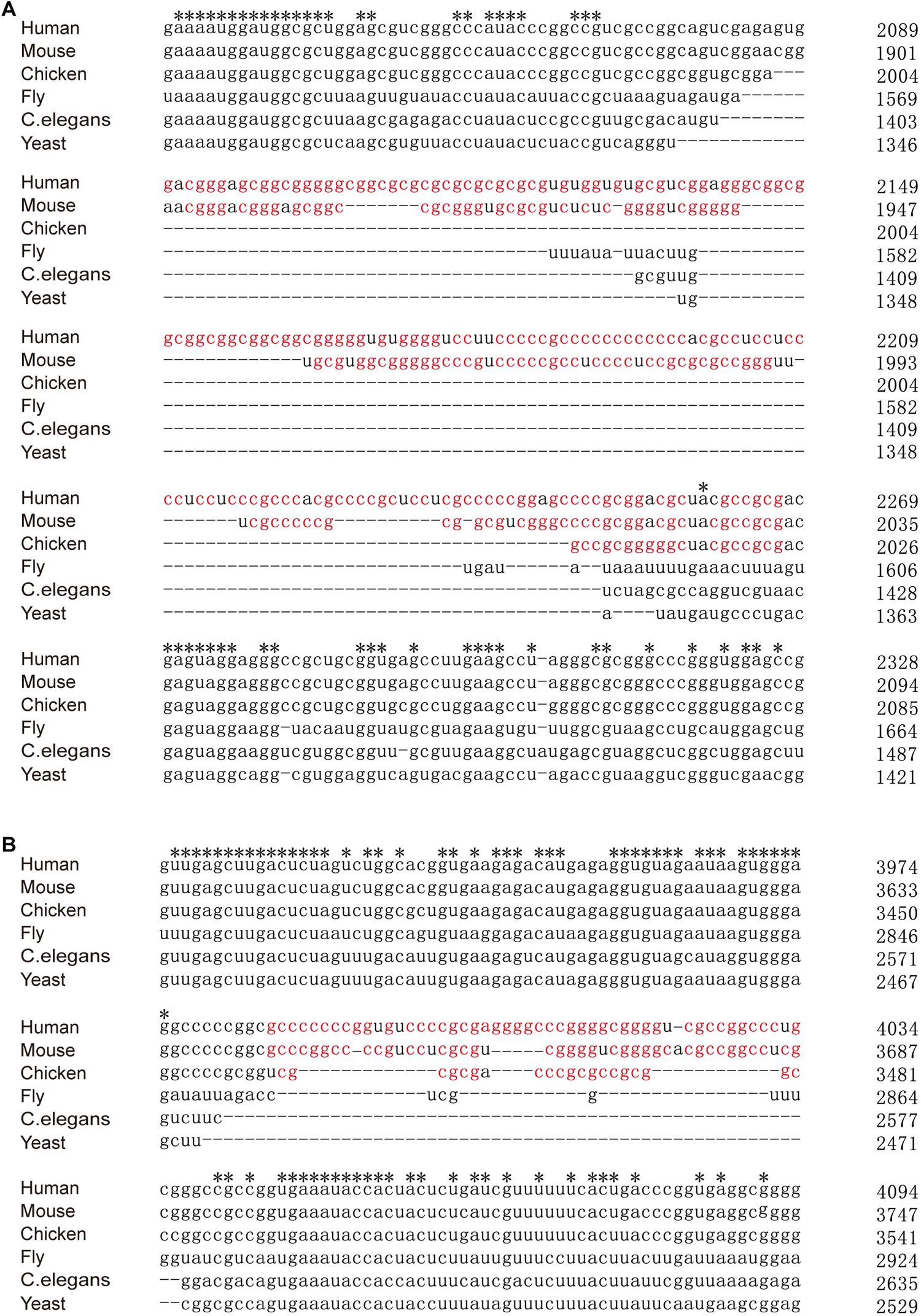
The extended sequences in 28S rRNA of higher organisms tend to be GC-rich. (**A**) Multiple sequence alignment of 28S rRNAs of the indicated organisms. Shown is a fragment of aligned 28S rRNAs with extended sequences in the higher organisms flanked by conserved sequences. Stars indicate identical nucleotides. G and C are marked red within the extended sequences. (**B**) as in (A), but another fragment of 28S rRNA.

**Figure S9.**
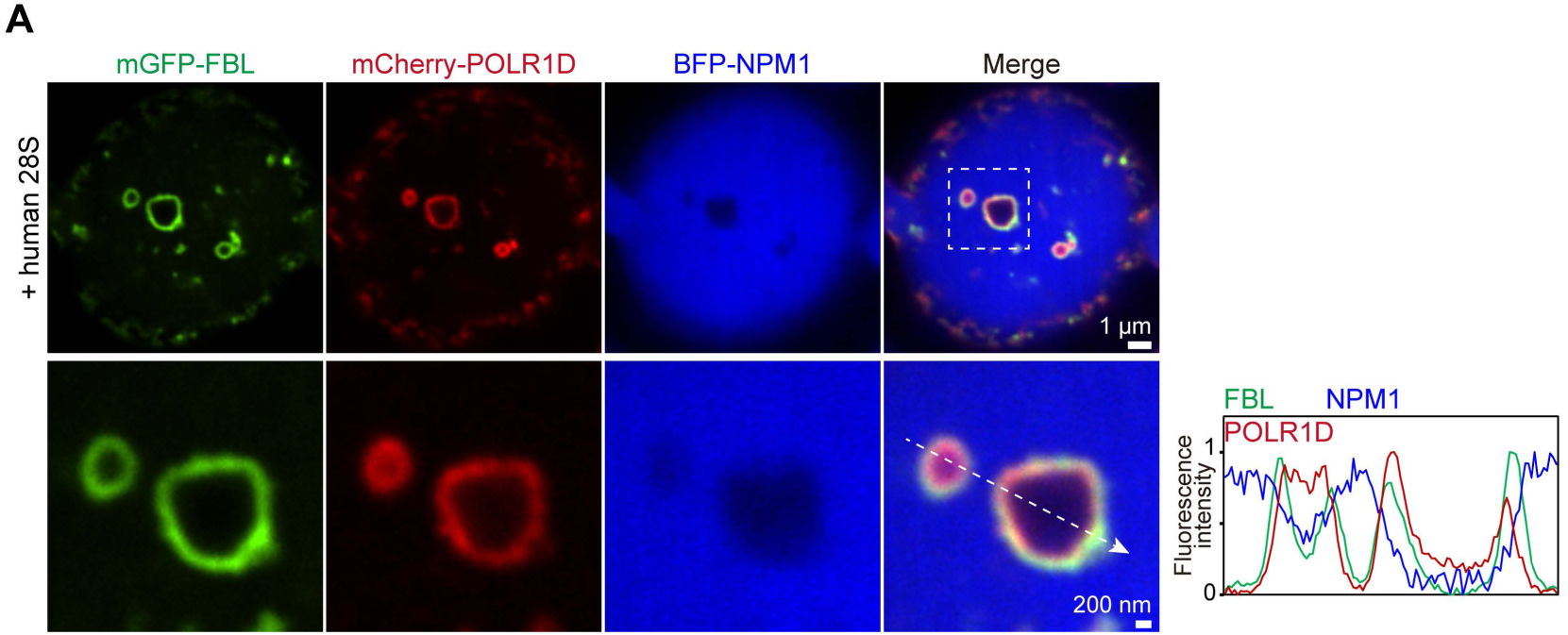
In the DFC-like structure with a large size, POLR1D localizes close to the rim of the FBL ring instead of filling the FBL shell. (**A**) Representative confocal images of phase separation experiments using purified mGFP-FBL (5 µM), BFP-NPM1 (30 µM), and mCherry-POLR1D (30 µM), in the presence of human 28S rRNA (2 µg/µl). Bottom right: line profile of fluorescence intensities. The arrow indicates the plane used for the line profile generation.

## Supplementary Files

**Supplementary File 1 primers used in the molecular cloning**

**Supplementary File 2 Sequence of 5.8S rRNAs**

**Supplementary File 3 Sequence of 18S rRNAs**

**Supplementary File 4 Sequence of 28S rRNAs**

## STAR Methods

### Cell lines

The human cervical cancer cell line, HeLa, was a gift from the lab of Christine Mayr (Memorial Sloan Kettering Cancer Center). The cell line has not been authenticated.

The HEK293T/17, U2OS, A549, and NIH/3T3, authenticated by STR profiling, were purchased from Procell. Cells were maintained at 37°C with 5% CO2 in Dulbecco’s Modified Eagle Medium containing 4500 mg/l glucose, 10% fetal bovine serum, 100 U/ml penicillin, and 100 mg/ml streptomycin. Mycoplasma detection was performed by DAPI staining.

### Constructs

To make pcDNA-puro-mGFP-FBL, the FBL coding sequence (CDS) was PCR-amplified from cDNA of HEK293T cells with primers FBL F and FBL R. FBL was inserted into the pcDNA3.1-puro-mGFP vector with HindIII and EcoRI restriction sites using T4 DNA ligase. FBL CDS was subcloned into pLVX-blast-mNeonGreen to make pLVX-blast-mNeonGreen-NPM1.

To make pcDNA-puro-BFP-NPM1, the NPM1 CDS was PCR-amplified from cDNA of HEK293T cells with primers NPM1 F and NPM1 R. NPM1 was inserted into the pcDNA3.1-puro-BFP (mTagBFP2) vector with HindIII and EcoRI restriction sites using T4 DNA ligase. NPM1 CDS was subcloned into pLVX-blast-mNeonGreen to make pLVX-blast-mNeonGreen-NPM1.

To make pcDNA-puro-mCherry-POLR1D, the POLR1D CDS was PCR-amplified from cDNA of HEK293T cells with primers POLR1D F and POLR1D R. POLR1D was inserted into the pcDNA3.1-puro-mCherry vector with BsrGI and EcoRI restriction sites through recombinational cloning.

mGFP-FBL, BFP-NPM1, and mCherry-POLR1D were subcloned into a modified pRSFDuet-1 vector. All constructs have an N-terminus starting sequence of MKHHHHHHHENLYFQ, which was cleaved by TEV protease during purification. Due to the TEV cleavage site, all proteins start with GA or G residues at the N-terminus.

The following are the sequences of the fusion proteins used in this study:

mEGFP-FBL

GAMVSKGEELFTGVVPILVELDGDVNGHKFSVSGEGEGDATYGKLTLKFICTTGKLPVPWPTLVTTLTYGVQCFSRYPDHMKQHDFFKSAMPEGYVQERTIFFKDDGNYKTRAEVKFEGDTLVNRIELKGIDFKEDGNILGHKLEYNYNSHNVYIMADKQKNGIKVNFKIRHNIEDGSVQLADHYQQNTPIGDGPVLLPDNHYLSTQSKLSKDPNEKRDHMVLLEFVTAAGITLGMDELYKKLMKPGFSPRGGGFGGRGGFGDRGGRGGRGGFGGGRGRGGGFRGRGRGGGGGGGGGGGGGRGGGGFHSGGNRGRGRGGKRGNQSGKNVMVEPHRHEGVFICRGKEDALVTKNLVPGESVYGEKRVSISEGDDKIEYRAWNPFRSKLAAAILGGVDQIHIKPGAKVLYLGAASGTTVSHVSDIVGPDGLVYAVEFSHRSGRDLINLAKKRTNIIPVIEDARHPHKYRMLIAMVDVIFADVAQPDQTRIVALNAHTFLRNGGHFVISIKANCIDSTASAEAVFASEVKKMQQENMKPQEQLTLEPYERDHAVVVGVYRPPPKVKN*

BFP-NPM1

GAMVSKGEELIKENMHMKLYMEGTVDNHHFKCTSEGEGKPYEGTQTMRIKVVEGGPLPFAFDILATSFLYGSKTFINHTQGIPDFFKQSFPEGFTWERVTTYEDGGVLTATQDTSLQDGCLIYNVKIRGVNFTSNGPVMQKKTLGWEAFTETLYPADGGLEGRNDMALKLVGGSHLIANAKTTYRSKKPAKNLKMPGVYYVDYRLERIKEANNETYVEQHEVAVARYCDLPSKLGHKLNKLMEDSMDMDMSPLRPQNYLFGCELKADKDYHFKVDNDENEHQLSLRTVSLGAGAKDELHIVEAEAMNYEGSPIKVTLATLKMSVQPTVSLGGFEITPPVVLRLKCGSGPVHISGQHLVAVEEDAESEDEEEEDVKLLSISGKRSAPGGGSKVPQKKVKLAADEDDDDDDEEDDDEDDDDDDFDDEEAEEKAPVKKSIRDTPAKNAQKSNQNGKDSKPSSTPRSKGQESFKKQEKTPKTPKGPSSVEDIKAKMQASIEKGGSLPKVEAKFINYVKNCFRMTDQEAIQDLWQWRKSL*

mCherry-POLR1-D,

GMVSKGEEDNMAIIKEFMRFKVHMEGSVNGHEFEIEGEGEGRPYEGTQTAKLKVTKGGPLPFAWDILSPQFMYGSKAYVKHPADIPDYLKLSFPEGFKWERVMNFEDGGVVTVTQDSSLQDGEFIYKVKLRGTNFPSDGPVMQKKTMGWEASSERMYPEDGALKGEIKQRLKLKDGGHYDAEVKTTYKAKKPVQLPGAYNVNIKLDITSHNEDYTIVEQYERAEGRHSTGGMDELYKEEDQELERKISGLKTSMAEGERKTALEMVQAAGTDRHCVTFVLHEEDHTLGNSLRYMIMKNPEVEFCGYTTTHPSESKINLRIQTRGTLPAVEPFQRGLNELMNVCQHVLDKFEASIKDYKDQKASRNESTF*

To make pTOPO-T7-human 5.8S, 18S, 28S rDNA, the human 5.8S, 18S, and 28s rRNA sequence (rDNA) were PCR amplified from cDNA of HEK293T cells with primers 5.8S human F and 5.8S human R, 18S human F and 18S human R and 28S human F and 28S human R, respectively.

To make pTOPO-T7-mouse 28S rDNA, the mouse 28S rDNA was PCR amplified from cDNA of L929 cells with primers 28S mouse F and 28S mouse R. The C.elegans 26S rDNA was PCR amplified from cDNA of C.elegans with primers 26S C.elegans F and 26S C.elegans R. The T7-5.8S, T7-18S, and T7-28S rDNA were inserted into the pTOPO vector through TOPO cloning (Easy-Do).

To make pTOPO-T7-chicken 28S rDNA, two cloning steps were performed. Step1: a fragment of 1,373 bp at the 3’ end of chicken 28S rDNA sequence was PCR-amplified from the cDNA of chicken cells and inserted into the pTOPO vector through TOPO cloning (Easy-Do) with a XbaI restriction site on the 5′ end and a XhoI restriction site on the 3′ end. The primers were chicken 28S F-1 and chicken 28S R-1. Step2: the remaining sequence of chicken 28S rRNA was divided into four fragments (fragment1, 2, 3, and 4) and were PCR-amplified and inserted into the plasmid in step1 with XbaI digestion through recombinational cloning (TransGen Biotech). The primers were fragment1: chicken 28S F-2 and chicken 28S R-2; fragment2: 28S F-3 and chicken 28S R-3, fragment3: chicken 28S F-4 and chicken 28S R-4, and fragment4: chicken 28S F-5 and chicken 28S R-5. The T7 promoter sequence was added on the 5′ end of fragment1.

To make pTOPO-T7-fly 26S rDNA, 26S rDNA was PCR-amplified from the cDNA of Drosophila melanogaster cells and inserted into the pTOPO vector through TOPO cloning (Easy-Do). The primers were Fly 26S F and Fly 26S R.

To make pTOPO-T7-yeast 25S rDNA, 25S rDNA was PCR-amplified from the cDNA of the baker’s yeast (Saccharomyces cerevisiae) and inserted into the pTOPO vector through TOPO cloning (Easy-Do). The primers were Yeast 25S-T7 F and Yeast 25S-T7 R.

For all pTOPO-rRNA constructs, a T7 promoter (TAATACGACTCACTATAGGG) was incorporated into the forward primers used to amplify the DNA templates. A unique restriction site was incorporated into the reverse primers for linearizing the plasmids for in vitro T7 transcription experiments.

The primers used are listed in the Supplementary File. 1.

### Recombinant protein purification

Plasmids of mEGFP-FBL, mTagBFP2-NPM1, mCherry-POLR1-H, mCherry-POLR1-D, and mCherry-POLR1-C in the modified pRSFDuet-1 vector were individually transformed into *E. coli* BL21 (DE3) cells. After transformation, a single colony was inoculated into 5 mL LB supplemented with 50 mg/L kanamycin at 220 rpm, 37°C. After 5 h, the culture was diluted into 100 mL LB medium supplemented with 50 mg/L kanamycin at 220 rpm, 37°C. After overnight growth, the culture was diluted into 700 mL LB medium supplemented with 50 mg/L kanamycin at 220 rpm, 37°C. Turbidity was monitored at a wavelength of 600 nm, and upon reaching an optical density (OD600) of 0.6 - 0.8, IPTG was added to the LB medium at the concentration of 0.5 mM for the induction of protein expression. After incubation for 24 h at 220 rpm, 16°C, cells were harvested by centrifugation (4,000 g, 15 min, 4°C), resuspended in 40 mL Binding buffer (20 mM Tris pH 7.5, 600 mM NaCl, 2mM DTT, 10% glycerol) and then lysed by high-pressure homogenizer at 4°C, 1,300 bar. After centrifugation at 15,000 rpm for 30 min at 4°C, the supernatant was collected and then flowed slowly through the Ni Sepharose Excel column (GE healthcare) for binding proteins. The unbound proteins were washed out with binding buffer and washing buffer (20 mM Tris pH 7.5, 600 mM NaCl, 2mM DTT, 10% glycerol, 15 mM imidazole). Bound proteins were eluted with elution buffer (20 mM Tris pH 7.5, 600 mM NaCl, 2mM DTT, 10% glycerol, 500 mM imidazole). TEV protease and RNase A were added in protein elution to remove 7xHis tag and nucleic acids during overnight dialysis against 2L of binding buffer at 4°C. Sample purity and TEV cleavage were confirmed on a 10% acrylamide SDS-PAGE gel stained with Coomassie. The digested product was transferred again into the Ni Sepharose Excel column to remove the 7xHis tag fragment and TEV protease. Then the protein was further purified by the gel filtration chromatography (Superdex-75 increase 10/300GL; GE Healthcare) equilibrated with binding Buffer (20 mM Tris pH 7.5, 600 mM NaCl, 2mM DTT, 10% glycerol). Sample purities were confirmed on a 10% acrylamide SDS-PAGE gel stained with Coomassie, and nucleic acid contamination was checked by the absorption values at 260nm and 280nm.

### Establishment of *RNASEL* knockout cell line

To generate RNASEL knockout cells, the px458-puro (pSpCas9(BB)-2A-Puro) CRISPR/Cas9 vector was used. The guide RNA (gRNA) sequence (GAATTTGAGGCGAAAGACAAAGG) was cloned into the BbsI site to make px458-puro-RNASEL gRNA. HeLa cells were plated in a 6-well at the density of 4 x 10^5^. 1 µg px458-puro-RNASEL gRNA plasmid was transfected into HeLa cells. 24 hours after transfection, puromycin was added into the medium to a final concentration of 2 µg/ml to kill the un-transfected cells. Puromycin was removed after 24 hours of selection, and cells were grown to about 80 % confluency. Then, cells were diluted sequentially and plated into 96-well plates with one cell in 200 µl medium per well. Cells were grown in the 96-wells for 10 days, then single clones were selected and expanded to a 24-well and a 6-well subsequently. Cells were harvested for protein extraction and DNA extraction. Western blot was first performed to identify clones without RNase L expression. HeLa cell clones with no expression of RNase L (*RNASEL* knockout) were kept. The genomic sequence flanking the gRNA targeting sequence was PCR-amplified and cloned into the pTOPO vector. DNA sequencing was performed to confirm the indel mutations in the knockout cells.

### Transfections

Endofectin MAX (Genecopoeia) was used for all transfections.

### Imaging

For Fig. 1d, the microscopy used was Nikon confocal AX with NSPARC, using an SR Apo TIRF 100X oil, 1.49 NA objective. Images were taken under the NSPARC mode (super-resolution).

For other images, the microscopy used was the Olympus IXplore SpinSR microscope equipped with Yokogawa CSU-W1 SoRa (SoRa disk) and operated with cellSens Dimension software. The objective is Apo TIRF 60X oil, 1.5 NA. For super-resolution imaging, Z-stacks with a step size of 210 nm were acquired under the super-resolution mode with the SoRa disk. Deconvolution was performed using the constrained iterative method with the default settings: Modality, Laser Scanning Confocal or Spinning Disk Confocal; Algorithm: Advanced Maximum Likelihood; Iteratopms, 5. Images were exported using the Olympus OlyVIA software.

### RNA FISH

RNA FISH was performed as described previously^44^ with a few modifications. HeLa cells were plated on 4-chambered glass-bottom dishes at a density of 0.2×10^6^ to grow overnight. Cells were fixed with 2% formaldehyde at 37°C for 15 minutes and washed twice for 5 min with PBS. For RNA FISH in cells treated with RNase A, after fixation, cells were permeabilized in 0.1% Triton X-100 in PBS for 10 min at RT. 200 µl of RNase A in PBS at a 200 µg/µl concentration was added to each chamber and kept at 37 LJ for 2 h. Cells were washed twice quickly with PBS. PBS was then discarded, and 0.5 mL 70% ethanol was added. The dishes were kept at 4°C for 2 to 8 hours. The 70% ethanol was aspirated, 0.5 mL wash buffer (2x SSC, 10% formamide in RNase-free water) was added, and incubated at RT for 5 min. Hybridization mix was prepared by mixing 10% Dextran sulfate, 10% formamide, 2 x SSC, 2 mM ribonucleoside vanadyl complex (NEB), 200 μg/ml yeast tRNA (Sigma, 10109495001), 20nM rRNA probe. A 100 μl hybridization mix was added to each chamber and hybridized at 37°C overnight. After hybridization, cells on dishes were washed twice for 30 min (each time 15 min) with pre-warmed wash buffer (1 ml, 37°C) in the dark, followed by one quick wash with PBS, and kept in PBS for imaging. Imaging was taken using the Olympus IXplore SpinSR microscope equipped with Yokogawa CSU-W1 SoRa.

FISH probes (3’ Rox fluorescently labeled DNA oligos) were synthesized by GENEWIZ Life Sciences. The sequences of rRNA probes were previously reported.

5’ETS human: ACGACGTCACCACATCGATCACGAAGAG

5S rRNA human: AAGCCTACAGCACCCGGTATTCCCA

5.8S rRNA human: CATCGACGCACGAGCCGAGTGATCCAC

18S rRNA human: GAGGTTTCCCGTGTTGAGTCAAATTAAGCCGCA

28S rRNA human: CCTTGTGTCGAGGGCTGACTTTCAATAG

### In vitro RNA transcription

All rRNAs were in vitro transcribed using the T7 high-yield RNA transcription kit (Vazyme). In vitro transcription was performed in a 20 µl volume according to the manufacturer’s guidelines. DNA templates used for in vitro transcription were linearized pTOPO-T7-5.8/18/28S rDNA vectors. The transcription reaction was incubated for 4 h at 37 °C, followed by DNase I treatment for 30 min at 37 °C to remove template DNAs, then transcribed RNA was precipitated with LiCl for 2 h at -20 °C. RNAs were centrifuged at 13,000 g for 15 min, and the RNA pellets were washed with 70% ethanol twice. RNAs were dissolved in 20 mM Tris, pH 7.4, and run on agarose gels to evaluate the integrity and size of the RNA. RNA concentration was measured by Nano300. RNAs were stored at -20 °C.

To generate Cy3-labeled 28S rRNA, 0.4 µl of 2.5 mM Cy3-UTP (APExBIO) was added into 20 µl of in vitro transcription reaction.

### Western blot

Western blots were performed as described previously^44^. Imaging was captured on the Odyssey CLx imaging system (Li-Cor). The antibodies used are rabbit anti-β-Actin (ABclonal, AC026), mouse anti-RNase L (Santa Cruz, sc-74405), IRDye 680RD donkey anti-mouse IgG secondary antibody (Li-COR Biosciences, 926-68072), and IRDye 800CW donkey anti-rabbit IgG secondary antibody (Li-COR Biosciences, 926-32213).

### RNA interaction and native agarose gel electrophoresis

RNA interaction experiements were perfomed as described previously with a few modifications^35^. RNAs were diluted into a 3 µl buffer (20 mM Tris pH 7.4, 150 mM NaCl) to a final concentration of 1-2 µg/µl, RNAs were annealed at 95°C for 2 min, 0.1 C/sec to 25 ^0^C in a PCR machine and then kept at 37 °C for 1 h. RNA native agarose gel electrophoresis was performed as described previously.

### Drug treatment in cells

Doxorubicin (stock 10 mM in water) or CX-5461 (stock 5 mM in water) was diluted to the intermediate concentration in water and then directly added to HeLa (mNeonGreen-FBL) cells to a final concentration of 2.5 µM doxorubicin or 1 µM CX-5461. Images were taken 30 min after adding the drug.

### Lentivirus production, infection, and generation of stable cell lines

To make stable cell lines, the lentiviral system with pLVX plasmid was used. pLVS/pdR8.2/VSVG plasmids were transfected into HEK293T/17 cells for lentiviral packaging. Virus was harvested 48 h after transfection. 50 μl virus was used per 6-well to infect HeLa cells. 24 h after infection, puromycin or blasticidin was added to the medium. Cells were selected in the medium with puromycin or blasticidin for 10 days to make stable cell lines.

### In vitro phase separation

All phase separation assays were performed in 20 µl phase separation buffer A (150 mM NaCl, 20 mM Tris-Cl, pH 7.4, 0.5 mM DTT, 2.5% glycerol, 5% PEG-8000 (Sigma)).

PEG-8000 stock (20%, w/w) was made in 20 mM Tris pH 7.4.

mGFP-FBL, BFP-NPM1, and mCherry-POLR1D protein stock were in buffer B (600 mM NaCl, 20 mM Tris-Cl, pH 7.4, 2 mM DTT, 10% glycerol). The high concentration (600 mM) of NaCl inhibits phase separation of the stock protein.

The final concentration of mGFP-FBL, BFP-NPM1, and mCherry-POLR1D was 5 µM, 30 µM, and 1 µM, respectively.

For phase separation experiments with the addition of rRNAs, rRNAs (stored in buffer C (20 mM Tris pH 7.4)) were diluted and mixed in 5 µl buffer D (150 mM NaCl, 20 mM Tris pH 7.4) to a desired concentration. For doxorubicin treatment, rRNA in buffer D was added to the doxorubicin solution (in 20 mM Tris pH7.4) to get a concentration of 250 µM of doxorubicin. Mixed RNA was kept at 37 LJ for 2 h before setting up the phase separation experiment.

For mGFP-FBL and BFP-NPM1 experiments, protein stocks were centrifuged at 13,000 g for 2 min to remove small protein aggregates. The supernatant was transferred into a new Eppendorf tube. mGFP-FBL and BFP-NPM1 were mixed in 5 µl buffer B. 5 ul buffer C, and 5 µl of 20% PEG-8000 was added into the 5 µl rRNA in buffer D and mixed well. The 15 µl mix was then added to the 5 µl protein solution and mixed well. Thus, NaCl concentration was diluted to 150 mM, and the final concentration of PEG-8000 was diluted to 5 % to allow phase separation.

The mixture (20 µl) was then transferred into a 384-well glass-bottom microplate (Cellvis). The microplate was kept in the dark at 37 LJ for 1 to 2 h, followed by imaging of the condensates using confocal microscopy.

For mGFP-FBL BFP-NPM1, and mCherry-POLR1D experiments, mGFP-FBL BFP-NPM1, and mCherry-POLR1D protein stocks were mixed in 5 µl buffer B. The phase separation experiment was performed as described above.

For phase separation with Cy3-labeled 28S rRNA, 4 µg of Cy3-28S rRNA and 36 µg of un-labeled 28S rRNA were mixed in buffer D. The phase separation experiment was performed as described above. The total reaction volume is 20 µl. Thus, the final concentration of rRNA was 2 µg/ µl (0.2 µg/µl Cy3-labeled + 1.8 µg/µl un-labeled 28S rRNA).

### Line profile

Line profile analysis was performed to examine whether rRNAs are co-localized with DFC in the nucleolus. Line profiles were generated with FIJI (ImageJ). A straight line was drawn across two to four adjacent DFC structures within a single nucleolus in HeLa cells staby expressing mNeonGreen-FBL. For each cell, two straight lines were drawn in different directions. Fluorescence signals of mNeonGreen-FBL protein and the examined rRNAs along the straight line were calculated with the plot profile tool in FIJI. The Pearson’s correlation coefficient (r) of two fluorescence signals was calculated in Excel.

### Counting the number of droplets with DFC-like structures in the in vitro phase separation experiment

To calculate the percentage of condensates with the formation of DFC-like structures, the total condensate number was counted first with FIJI. Small condensates were excluded to avoid potential small protein aggregates. Condensates with a more than 1 µm diameter were selected and counted automatically by FIJI. The number of condensates with DFC-like structures was counted manually in each slice of the z-stack images.

### Quantification of the extent of high molecular weight rRNA complexes in native agarose gel

To quantify the degree of multivalency of the 28S rRNA from different species, RNA interaction, and native agarose gel electrophoresis experiments were performed. The agarose gel images were analyzed with FIJI. An ROI (region of interest) was first drawn to cover a whole lane, and this lane’s total intensity was calculated (intensity-total). Then, a second ROI was drawn to cover the high molecular weight region (above the main band), and the intensity was calculated (intensity-high). The ratio of intensity-high / intensity-total indicates the degree of the multivalency of each 28S rRNA.

### Statistical analysis and reproducibility

All the histograms and X-Y plots were generated with GraphPad Prism. For the pair-wise comparisons, a two-tailed Student’s *t*-test was performed. NS means not significant; * means p<0.05; ** means p<0.01; *** means p<0.001. Error bars represent means ± SD.

For all the figure data, at least three independent experiments were performed.

