## Supplementary material for "Multivalent 28S rRNA Is the Organizer of the Nucleolus’s Multi-layered Architecture": Table S1

```
CLUSTAL O(1.2.4) multiple sequence alignment


Human           cgacucuuagcgguggaucacucggcucgugcgucgaugaagaacgcagcuagcugcgag      60
Mouse           -gacucuuagcgguggaucacucggcucgugcgucgaugaagaacgcagcuagcugcgag      59
Chicken         caacucuuagcgguggaucacucggcucgugcgucgaugaagaacgcagcuagcugcgag      60
Fly             -aacucuaagcgguggaucacucggcucaugggucgaugaagaacgcagcaaacugugcg      59
C.elegans       cuagcuucagcgauggaucgguugcaucgaguaucgaugaagaacgcagcuugcugcguu      60
Yeast           aaacuuucaacaacggaucucuugguucucgcaucgaugaagaacgcagcgaaaugcgau      60
                  *   * * *   *****  * *  **  *  ****************     ** *

Human           aauuaaugugaauugcaggacacauu-gaucaucgacacuucgaacgcacuug-cggccc      118
Mouse           aauuaaugugaauugcaggacacauu-gaucaucgacacuucgaacgcacuug-cggccc      117
Chicken         aauuaaugugaauugcaggacacauu-gaucaucgacacuucgaacgcacuug-cggccc      118
Fly             ucaucgugugaacugcaggacacau--gaacaucgacauuuugaacgcauaucgcagucc      117
C.elegans       acuuaccacgaauugcagacg-c-uuagaguggugaaauuucgaacgcauag--caccaa      116
Yeast           acguaaugugaauugcagaauuccgugaaucaucgaaucuuugaacgcacauugcgcccc      120
                   *     *** *****    *     *     **   ** *******     *

Human           cggguuccucccggggcuacgccugucugagcgucgcuu-   157
Mouse           cggguuccucccggggcuacgccugucugagcgucgguug   157
Chicken         cggguuccucccggggcuacgccugccugagcgucgcuu-   157
Fly             augcu-------g---------------------------   123
C.elegans       cug-ggccuccaguugguacgucugguucaggguuguu--   153
Yeast           uug-guauuccagggggcaugccuguuugagcgucauuu-   158
                  *         *
```
