## Supplementary material for "Multivalent 28S rRNA Is the Organizer of the Nucleolus’s Multi-layered Architecture": Table S2

CLUSTAL O(1.2.4) multiple sequence alignment

```
Human        -uaccugguugauccugccaguag-cauaugcuugucucaaagauuaagccaugcauguc      58
Mouse        -uaccugguugauccugccaguag-cauaugcuugucucaaagauuaagccaugcauguc      58
Chicken      -uaccugguugauccugccaguag-cauaugcuugucucaaagauuaagccaugcauguc      58
Fly          -auucugguugauccugccaguaguuauaugcuugucucaaagauuaagccaugcauguc      59
C.elegans    auaccugauugauucugucagcgc-gauaugcucaaguaaaagauuaagccaugcaugcu      59
Yeast        -uaucugguugauccugccaguagucauaugcuugucucaaagauuaagccaugcauguc      59
                 *** ***** *** ***     *******    * *******************

Human        uaaguacgcacggccgguacagugaaacugcgaauggcucauuaaaucaguuaugguucc      118
Mouse        uaaguacgcacggccgguacagugaaacugcgaauggcucauuaaaucaguuaugguucc      118
Chicken      uaaguacacacgggcgguacagugaaacugcgaauggcucauuaaaucaguuaugguucc      118
Fly          uaaguacacacgaauua-aaagugaaaccgcaaaaggcucauuauaucaguuaugguucc      118
C.elegans    uugauu----------caucaaugaaauugcguacggcucauuagagcagauaucaccu-      108
Yeast        uaaguauaagcaauuuauacagugaaacugcgaauggcucauuaaaucaguuaucguuua      119
             *   *               * *****  **  * ********* * *** ***

Human        uuuggucgcucgcuccucuccuacu-uggauaacugugguaauucuagagcuaauacaug      177
Mouse        uuuggucgcucgcuccucuccuacu-uggauaacugugguaauucuagagcuaauacaug      177
Chicken      uuuggucgcuccccuc-ccguuacu-uggauaacugugguaauucuagagcuaauacaug      176
Fly          uuagaucguuaaca-----guuacu-uggauaacugugguaauucuagagcuaauacaug      172
C.elegans    -ua--uc------cgggauccucauauggauaacugcggaaauacuggagcuaauacaug      159
Yeast        uuugauaguuccu----uuacuacaugguauaacugugguaauucuagagcuaauacaug      175
              *   *               *     * ******* ** *** ** *************

Human        ccgacgggcgcugacccccuu-cg-cggggggaugcgugcauuuaucagaucaaaacca      235
Mouse        ccgacgggcgcugacccccuucccggggggggaugcgugcauuuaucagaucaaaacca      237
Chicken      ccgacgagcgccgaccuc----------cggggacgcgugcauuuaucagaccaaaacca      226
Fly          caauuaaaacaugaa---ccuu------augggacaugugcuuuuauuaggcuaaaacca      223
C.elegans    caacuauaccccaacg------------caaggcggggugcaauuauuagaacagacca-      206
Yeast        cuuaaaaucucgac----ccu-------uuggaagagauguauuuauuagauaaaaaauc      224
             *                              *      **   **** **   * *

Human        acccggucagccccucuccggccccggccggggggcgggcgccggcggcuuuggu-gacu      294
Mouse        acccggugagcucccucccggcuccggccgggggucgggcgccggcggcuuuggu-gacu      296
Chicken      acccgggcuc-----------------------------gcccggcggcuuuggu-gacu      256
Fly          agc----gaucgcaa------------------------gaucguuauauugguugaacu      255
C.elegans    aac----guuuc------------------------ggacgu-----uguuuguu-gacu      233
Yeast        aau----gu------------------------------cuucggacucuuugau-gauu      249
             *                                                ** * *  * *

Human        cuagauaaccucgggccgaucgcacgcccc--ccguggcggcgacgacccauucgaacgu      352
Mouse        cuagauaaccucgggccgaucgcacgcccc--ccguggcggcgacgacccauucgaacgu      354
Chicken      cuagauaaccucgagccgaucgcacgccc---ccguggcggcgacgacccauucgaaugu      313
Fly          cuagauaacaug---cagaucguaugguc---uuguaccgacgacagaucuuucaaaugu      309
C.elegans    cugaauaaagca-----guuu--acugucaguuucgacugacucuauccggaaagggugu      286
Yeast        cauaauaacuuuu--cgaaucgcauggcc---uugugcuggcgaugguucauucaaauuu      304
             *   ****           *   *    *          * *                 *

Human        cugcccuaucaacuuucgaugguagucgccgugccuaccauggugaccacgggugacggg      412
Mouse        cugcccuaucaacuuucgaugguagucgccgugccuaccauggugaccacgggugacggg      414
Chicken      cugcccuaucaacuuucgaugguacugucugugccuaccauggugaccacggguaacggg      373
Fly          cugcccuaucaacuuuugaugguaguaucuaggacuaccaugguugcaacggguaacggg      369
C.elegans    cugcccuuucaacu--agaugguaguuuauuggacuaccaugguuguuacggguaacgga      344
Yeast        cugcccuaucaacuuucgaugguaggauaguggccuaccaugguuucaacggguaacggg      364
             ******* ******   *******        * **********    ****** ****

Human        gaaucaggguucgauuccggagagggagccugagaaacggcuaccacauccaaggaaggc      472
Mouse        gaaucaggguucgauuccggagagggagccugagaaacggcuaccacauccaaggaaggc      474
Chicken      gaaucaggguucgauuccggagagggagccugagaaacggcuaccacauccaaggaaggc      433
Fly          gaaucaggguucgauuccggagagggagccugagaaacggcuaccacaucuaaggaaggc      429
C.elegans    gaauaaggguucgacuccggagagggagccuuagaaacggcuaccacguccaaggaaggc      404
Yeast        gaauaaggguucgauuccggagagggagccugagaaacggcuaccacauccaaggaaggc      424
             **** ********* **************** *************** ** *********
```

```
Human       agcaggcgcgcaaauuacccacucccgacccggggagguagugacgaaaaauaacaauac    532
Mouse       agcaggcgcgcaaauuacccacucccgacccggggagguagugacgaaaaauaacaauac    534
Chicken     agcaggcgcgcaaauuacccacucccgacccggggagguagugacgaaaaauaacaauac    493
Fly         agcaggcgcguaaauuacccacucccagcucggggagguagugacgaaaaauaacaauac    489
C.elegans   agcaggcgcgaaacuuauccacuguugagua--ugagauagugacuaaaaauauaaaga-    461
Yeast       agcaggcgcgcaaauuacccaauccuaauucagggagguagugacaauaaauaacgauac    484
            ********** ** *** *** *           *** ******* * *****   * *

Human       aggacucu--uucgaggcccuguaauuggaaugaguccacuuuaaauccuuuaacgagga    590
Mouse       aggacucu--uucgaggcccuguaauuggaaugaguccacuuuaaauccuuuaacgagga    592
Chicken     aggacucu--uucgaggcccuguaauuggaaugaguccacuuuaaauccuuuaacgagga    551
Fly         aggacucauauccgaggcccuguaauuggaaugaguacacuuuaaauccuuuaacaagga    549
C.elegans   ----cucauccuuuuggaugaguuauuucaaugaguugaauacaaaugauucuucgagua    517
Yeast       agggccca--uucgg-gucuuguaauuggaaugaguacaauguaaauaccuuaacgagga    541
                * *         *    ** ***  *******  * *  ****   *   * ** *

Human       uccauuggagggcaagucuggugccagcagccgcgguaauuccagcuccaauagcguaua    650
Mouse       uccauuggagggcaagucuggugccagcagccgcgguaauuccagcuccaauagcguaua    652
Chicken     uccauuggagggcaagucuggugccagcagccgcgguaauuccagcuccaauagcguaua    611
Fly         ccaauuggagggcaagucuggugccagcagccgcgguaauuccagcuccaauagcguaua    609
C.elegans   gcaaggagagggcaagucuggugccagcagccgcgguaauuccagcucuccuaguguauc    577
Yeast       acaauuggagggcaagucuggugccagcagccgcgguaauuccagcuccaauagcguaua    601
             * *   *****************************************   *** ****

Human       uuaaaguugcugcaguuaaaaagcucguaguuggaucuugggagcgggcgggcgguccgc    710
Mouse       uuaaaguugcugcaguuaaaaagcucguaguuggaucuugggagcgggcgggcgguccgc    712
Chicken     uuaaaguugcugcaguuaaaaagcucguaguuggaucuugggaucgagcuggcgguccgc    671
Fly         uuaaaguuguugcgguuaaaacguucguaguugaacuugugcuucauacggguaguacaa    669
C.elegans   ucguuauugcugcgguuaaaaagcucguaguuggaucuagguuacgugccgcaguuc---    634
Yeast       uuaaaguuguugcaguuaaaaagcucguaguugaacuuugggcccgguuggccgguccga    661
            *     *** *** ******* * ********* *  *  *   *     *    *

Human       cgc--------------------------------gaggcgagccaccgcc---cgucc    734
Mouse       cgc--------------------------------gaggcgagucaccgcc---cgucc    736
Chicken     cgc--------------------------------gaggcgagcuaccgcc---ugucc    695
Fly         cuuacaauugugguuaguacuauaccuuuauguauguaagcguauuaccgguggaguucu    729
C.elegans   ---gcaauuu-----------------------------gcg--ucaacuguggu-----    655
Yeast       uuu--------------------------------uu-ucgu-guacugga---uuucc    683
                                                    **    *

Human       --c---------------cgccccuu---------gccucucg---gcgcc---cccucg    762
Mouse       --c---------------cgccccuu---------gccucucg---gcgcc---cccucg    764
Chicken     --c---------------agcccc-u---------gucucucg---gcgcc---cccucg    722
Fly         uauaugugauuaaauacuuguaucuuuucauauguuccuccuauuuaaaaaccugcauua    789
C.elegans   ------------------cgugacuucuaauuugcugg-----uuuga------gguugg    686
Yeast       aac---------------ggggccuu---------uccuucug---gcuaa---ccuuga    713
                               *   * *                               *

Human       augcucuuagcugagugucccgcgg-ggcccgaagcguuuacuuugaaaaaauuagagug    821
Mouse       augcucuuagcugagugucccgcgg-ggcccgaagcguuuacuuugaaaaaauuagagug    823
Chicken     augcucuuaacugagugucccgcgg-ggcccgaagcguuuacuuugaaaaaauuagagug    781
Fly         gugcucuuaaacgaguguua--uugugggccgguacuauuacuuugaacaaauuagagug    847
C.elegans   guu-------------cgcccuucaacugccagcagguuuaccuugaauaaaucagagug    733
Yeast       guccuugu--------ggcucuuggcgaaccaggacuuuuacuuugaaaaaauuagagug    765
             *                           **       **** ***** **** ******

Human       uucaaagcaggcccgagccgccuggauacc-gcagcuaggaauaauggaauaggacc-gc    879
Mouse       uucaaagcaggcccgagccgccuggauacc-gcagcuaggaauaauggaauaggacc-gc    881
Chicken     uucaaagcaggcu--ggccgccggaauacu-ccagcuaggaauaauggaauaggacu-cc    837
Fly         cuuaaagcaggcuucaaaugccugaauauucugugcaugggauaaugaaauaagaccucu    907
C.elegans   cucaauacaagcg---cuugcuugaauagc-ucaucauggaauaaugaaacaggacuucg    789
Yeast       uucaaagcaggcg--uauugcucgaauaua-uuagcauggaauaauagaauaggacguuu    822
             * **  ** **       **  * ***       *  ** *****  ** * ***
```

```
Human       gguucuauuuuguugguuuucggaacu-gaggccaugauuaagagggacgg-ccgggggc     937
Mouse       gguucuauuuuguugguuuucggaacu-gaggccaugauuaagagggacgg-ccgggggc     939
Chicken     gguucuauuuuguugguuuucggaaac-ggggccaugauuaagagggacgg-ccgggggc     895
Fly         guucugcuuucauugguuuucagaucaagagguaaugauuaauagaagcaguuugggggc     967
C.elegans   guucu--uuuuguuggu-ucuagaacu-gauuuaaugguuaagagggacaaaccgggggc     845
Yeast       gguucuauuuuguugguuucuaggacc-aucguaaugauuaauagggacgg-ucgggggc     880
            * *    ***  ***** *   *           *** **** **   *     ******

Human       auucguauugcgccgcuagaggugaaauucuuggaccggcgcaagacggaccagagcgaa     997
Mouse       auucguauugcgccgcuagaggugaaauucuuggaccggcgcaagacggaccagagcgaa     999
Chicken     auucguauugugccgcuagaggugaaauucuuggaccggcgcaagacgaacuaaagcgaa     955
Fly         auuaguauuacgacgcgagaggugaaauucuuggaccgucguaagacuaacuuaagcgaa     1027
C.elegans   auucguaucauuacgcgagaggugaaauucguggaccguagugagacgcccaacagcgaa     905
Yeast       aucaguauucaauugucagaggugaaauucuuggauuuauugaagacuaacuacugcgaa     940
            **  ****      *  ************* ****        ****   *    *****

Human       agcauuugccaagaauguuuucauuaaucaagaacgaaagucggagguucgaagacgauc     1057
Mouse       agcauuugccaagaauguuuucauuaaucaagaacgaaagucggagguucgaagacgauc     1059
Chicken     agcauuugccaagaauguuuucauuaaucaagaacgaaagucggagguucgaagacgauc     1015
Fly         agcauuugccaaagauguuuucauuaaucaagaacgaaaguuagagguucgaaggcgauc     1087
C.elegans   agcauuugccaagaaugucuucauuaaucaagaacgaaagucagagguucgaaggcgauu     965
Yeast       agcauuugccaaggacguuuucauuaaucaagaacgaaaguuaggggaucgaagaugauc     1000
            ************  * ** **********************  * ** ******  ***

Human       agauaccgucguaguuccgaccauaaacgaugccgaccggcgaugcggcgguuauucc     1117
Mouse       agauaccgucguaguuccgaccauaaacgaugccgacuggcgaugcggcgguuauucc     1119
Chicken     agauaccgucguaguuccgaccauaaacgaugccgacucgcgauccggcgguuauucc     1075
Fly         agauaccgcccuaguucuaaccauaaacgaugccagcuagcaauuggguguagcuacuuu     1147
C.elegans   agauaccgcccuaguucugaccguaaacgaugccaucucgcgauucggaggguuu-----     1020
Yeast       agauaccgucguagucuuaaccauaaacuaugccgacuagggaucggguggugguuuuuu     1060
            ******** * ****    *** ***** *****  *  *  **  ** *    *

Human       caugacccgccgggcagcuu-ccgggaaaccaaagucuuugguuccgggggaguaugg     1176
Mouse       caugacccgccgggcagcuu-ccgggaaaccaaagucuuugguuccgggggaguaugg     1178
Chicken     caugacccgccgggcagcuc-ccgggaaacccaagucuuugguuccgggggaguaugg     1134
Fly         uauggcucucucagucgcuucccgggaaaccaaagcuuuugggcuccgggggaaguaugg     1207
C.elegans   -uugcccugccgaggagcua-uccggaaacgaaagucuuucgguuccgggggguaguaugg     1078
Yeast       aaugacccacucggcaccuu-acgagaaaucaaagucuuugguucugggggggaguaugg     1119
              ** *   *   *   **   *  ****   ***  *** ** ** ***** *******

Human       uugcaaagcugaaacuuaaaggaauugacggaagggcaccaccaggaguggagccugcgg     1236
Mouse       uugcaaagcugaaacuuaaaggaauugacggaagggcaccaccaggaguggagccugcgg     1238
Chicken     uugcaaagcugaaacuuaaaggaauugacggaagggcaccaccaggaguggagccugcgg     1194
Fly         uugcaaagcugaaacuuaaaggaauugacggaagggcaccaccaggaguggagccugcgg     1267
C.elegans   uugcaaagcugaaacuuaaagaaauugacggaagggcaccacaaggcguggagcuugcgg     1138
Yeast       ucgcaaggcugaaacuuaaaggaauugacggaagggcaccaccaggaguggagccugcgg     1179
            * **** ************** ******************** *** ******* *****

Human       cuuaauuugacucaacacgggaaaccucacccggcccggacacggacaggauugacagau     1296
Mouse       cuuaauuugacucaacacgggaaaccucacccggcccggacacggacaggauugacagau     1298
Chicken     cuuaauuugacucaacacgggaaaccucacccggcccggacacggacaggauugacagau     1254
Fly         cuuaauuugacucaacacgggaaaacuuaccagguccgaacauaaguguguaagacagau     1327
C.elegans   cuuaauuugacucaacacgggaaaacucacccgguccggacaccauuaggacugacagau     1198
Yeast       cuuaauuugacucaacacggggaaacucaccagguccagacacaauaaggauugacagau     1239
            ********************* ** ** *** ** **  ***       *   *******

Human       ugauagcucuuucucgauuccgugggugguggugcauggccguucuuaguugguggagcg     1356
Mouse       ugauagcucuuucucgauuccgugggugguggugcauggccguucuuaguugguggagcg     1358
Chicken     ugagagcucuuucucgauuccgugggugguggugcauggccguucuuaguugguggagcg     1314
Fly         ugauagcucuuucucgaaucuaugggugguggugcauggccguucuuaguucguggagug     1387
C.elegans   ugaaagcucuuucucgauuuggugguugguggugcauggccguucuuaguugguggagug     1258
Yeast       ugagagcucuuucuugauuuugugggugguggugcauggccguucuuaguugguggagug     1299
            *** ********** ** *   *** ************************* ****** *
```

```
Human       auuugucugguuaauuccgauaacgaacgagacucuggcaugcuaacuaguuacgcga--    1414
Mouse       auuugucugguuaauuccgauaacgaacgagacucuggcaugcuaacuaguuacgcga--    1416
Chicken     auuugucugguuaauuccgauaacgaacgagacucuggcaugcuaacuaguuacgcga--    1372
Fly         auuugucugguuaauuccgauaacgaacgagacucaaauauuaaauagauaucuucag    1447
C.elegans   auuugucugguuuauuccgauaacgagcgagacucuagccugcuaaauaguuggcgaa--    1316
Yeast       auuugucugcuuaauugcgauaacgaacgagaccuuaaccuacuaaauaguggugcua--    1357
            ********* ** *** ********* ******       *  *** ***

Human       ------------------------ccc---------------------------------    1417
Mouse       ------------------------ccc---------------------------------    1419
Chicken     ------------------------ccc---------------------------------    1375
Fly         gauuauggugcugaagcuuauguagccuucauucauguuggcaguaaaaugcuuauugug    1507
C.elegans   -------------------------uc---------------------------------    1318
Yeast       ------------------------gca---------------------------------    1360

Human       --------------------------ccgagcggucggc--------------guccccc    1437
Mouse       --------------------------ccgagcggucggc--------------guccccc    1439
Chicken     --------------------------ccgagcggucggc--------------guc---c    1392
Fly         uuugaauguguuuauguaaguggagccguaccuguugguuugucccauuauaaggacacu    1567
C.elegans   ---------------------------------uucggg--------------uucguau    1331
Yeast       ---------------------------------uuugcu--------------gguuauc    1373
                                              * *

Human       aacuucuuagagggacaaguggcguucagccacccga-gauuga-gcaauaacaggucug    1495
Mouse       aacuucuuagagggacaaguggcguucagccacccga-gauuga-gcaauaacaggucug    1497
Chicken     aacuucuuagagggacaaguggcguucagccacccga-gauuga-gcaauaacaggucug    1450
Fly         agcuucuuaaauggacaaauugcgucuagcaauaaugagauuga-gcaauaacaggucug    1626
C.elegans   aacuucuuagagggauaagcgguguuuagccgcacga-gauuga-gcgauaacaggucug    1389
Yeast       cacuucuuagagggacuaucgguuucaagccgauggaaguuugaggcaauaacaggucug    1433
              ******* * ***  *   *  *  ***        * **** ** ************

Human       ugaugcccuuagauguccggggcugcacgcgcgcuacacugacuggcucagcgugugccu    1555
Mouse       ugaugcccuuagauguccggggcugcacgcgcgcuacacugacuggcucagcgugugccu    1557
Chicken     ugaugcccuuagauguccggggcugcacgcgcgcuacacugacuggcucagcuugugucu    1510
Fly         ugaugcccuuagauguccugggcugcacgcgcgcuacaaugaaaguaucaacgu--guau    1684
C.elegans   ugaugcccuuagauguccggggcugcacgcgugcuacacugguggagucagcgg--guuu    1447
Yeast       ugaugcccuuagacguucugggccgcacgcgcgcuacacugacggagccagcga--gucu    1491
            ************* ** * **** ******* ****** **   *   ** *    *  *

Human       acccuacgccggcaggcgcggguaacccguugaaccccauucgugauggggaucggggau    1615
Mouse       acccuacgccggcaggcgcggguaacccguugaaccccauucgugauggggaucggggau    1617
Chicken     acccuacgccggcaggcgcggguaacccguugaaccccauucgugauggggaucggggau    1570
Fly         uuccuagaccgagagguccggguaaaccgcugaaccacuuucaugcuugggauugugaac    1744
C.elegans   uuccuaugccgaaagguaucgguaaaccguugaaauucuuccauguccgggauaggguau    1507
Yeast       aaccuuggccgagaggucuugguaaucuugugaaacuccgucgugcuggggauagagcau    1551
              ***   ***  ***    ***** *   ****   *   * **   ***** * * *

Human       ugcaauuauuccccaugaacgaggaauucccaguaagugcgggucauaagcuugcguuga    1675
Mouse       ugcaauuauuccccaugaacgaggaauucccaguaagugcgggucauaagcuugcguuga    1677
Chicken     ugcaauuauuccccaugaacgaggaauucccaguaagugcgggucauaagcucgcguuga    1630
Fly         ugaaacuguu-cacaugaacuuggaauucccaguaagugugagucauuaacucgcauuga    1803
C.elegans   uguaauuauugcccuuaaacgaggaaugccuaguaagugugagucaucagcucacguuga    1567
Yeast       uguaauuauugcucuucaacgaggaauuccuaguaagcgcaagucaucagcuugcguuga    1611
            ** ** * ** * * * ***  ***** ** ****** *   ***** * **  * ****

Human       uuaagucccugcccuuuguacacaccgcccgucgcuacuaccgauuggaugguuuaguga    1735
Mouse       uuaagucccugcccuuuguacacaccgcccgucgcuacuaccgauuggaugguuuaguga    1737
Chicken     uuaagucccugcccuuuguacacaccgcccgucgcuacuaccgauuggaugguuuaguga    1690
Fly         uuacgucccugcccuuuguacacaccgcccgucgcuacuaccgauugaauuauuuaguga    1863
C.elegans   uuacgucccugcccuuuguacacaccgcccgucgcuauccgggacugaacugauucgaga    1627
Yeast       uuacgucccugcccuuuguacacaccgcccgucgcuaguaccgauugaauggcuuaguga    1671
            *** *********************************     ** ** *    ** * **
```

```
Human        ggcccucggaucggccccgccggggucggcccacggcccuggcggagcgcugagaag---    1792
Mouse        ggcccucggaucggccccgccggggucggcccacggcccuggcggagcgcugagaag---    1794
Chicken      gguccucggaucggccccggcggggucggc-cacggcccugccggagcgucgagaag---    1746
Fly          ggucuccggacgugaucacugugacgccuugcguguuacggu--uguuucgcaaaag---    1918
C.elegans    agaguggggacugucgcuucgagguu--uaacgacuuc--gu--uguugcggaaaccauu    1681
Yeast        ggccucaggaucugcuuagagaagg---gggcaacuccaucu--cagagcggagaau---    1723
              *     ***                     *                    * *

Human        acggucgaacuugacuaucuagaggaaguaaaagucguaacaagguuuccguaggugaac    1852
Mouse        acggucgaacuugacuaucuagaggaaguaaaagucguaacaagguuuccguaggugaac    1854
Chicken      acggucgaacuugacuaucuagaggaaguaaaagucguaacaagguuuccguaggugaac    1806
Fly          uugaccgaacuugauuauuuagaggaaguaaaagucguaacaagguuuccguaggugaac    1978
C.elegans    uuuaucgcauu----gguuugaaccggguaaaagucguaacaagguagcuguaggugaac    1737
Yeast        uuggacaaacuuggucauuuagaggaacuaaaagucguaacaagguuuccguaggugaac    1783
                  *  * *      * *  *     *****************   * **********

Human        cugcggaaggaucauua-                                              1869
Mouse        cugcggaaggaucauuaa                                              1872
Chicken      cugcggaaggaucauua-                                              1823
Fly          cugcggaaggaucauua-                                              1995
C.elegans    cugcagcuggaucaucg-                                              1754
Yeast        cugcggaaggaucauua-                                              1800
             **** *  *******
```
