## Supplementary material for "Multivalent 28S rRNA Is the Organizer of the Nucleolus’s Multi-layered Architecture": Table S3

CLUSTAL O(1.2.4) multiple sequence alignment

```
Human         -----cgcgaccucagaucagacguggcgacccgcugaauuuaagcauauuagucagcgg   55
Mouse         -----cgcgaccucagaucagacguggcgacccgcugaauuuaagcauauuagucagcgg   55
Chicken       -----cgcgaccucaggucagacguggcgacccgcugaauuuaagcauauuagucagcgg   55
Fly           uuauauacaaccucaacucauaugggacuaccccugaauuuaagcauauuaauuagggg   60
C.elegans     -----cucaaccugaacucagucgugauuacccgcugaacuuaagcauaucauuuagcgg   55
Yeast         -----uuugaccucaaaucagguaggaguacccgcugaacuuaagcauaucaauaagcgg   55
                       **** *  ***     *   **** ***** ********** * * ** **

Human         aggagaagaaacuaaccaggauucccucaguaacggcgagugaacagggaagagcccagc   115
Mouse         aggaaaagaaacuaaccaggauucccucaguaacggcgagugaacagggaagagcccagc   115
Chicken       aggaaaagaaacuaacgaggauucccucaguaacggcgagugaagagggaagagcccagc   115
Fly           aggaaaagaaacuaacaaggauuuucuuaguagcggcgagcgaaaagaaaacaguucagc   120
C.elegans     aggaaaagaaacuaaaaaggauucccuuaguaacggcgagugaaacgggaagagcccagc   115
Yeast         aggaaaagaaaccaaccgggauugccuuaguaacggcgagugaagcggcaaaagcucaaa   115
              **** ******* **   *****  ** **** ******* ***  *  ** **  **

Human         gccgaauccccgccccgc-ggcggggcgcgggacauguggcguacggaagacccgcuccc   174
Mouse         gccgaauccccgccgcgc-gucgcggcgugggaaauguggcguacggaagacccacuccc   174
Chicken       gccgaauccccgccccgc-gguggggcgcgggagguguggcguacggaagcccccauccc   174
Fly           acuaagucacuuugucua-uauggcaaaugugagaugcaguguauggagcgucaauauuc   179
C.elegans     gccgaaucg---aucagucuuuggcugcuucgaaauguggcguauaggug--uaaguuuc   170
Yeast         uuugaaaucugguaccuucggugcccgaguuguaauuuggag----aggg--caacuuug   169
                  *                 *        *   *   * *

Human         cggcgccgcucguggggggcccaaguccuucuga-ucgaggccc--agcccguggacggu   231
Mouse         cggcgccgcucguggggggcccaaguccuucuga-ucgaggccc--agcccguggacggu   231
Chicken       cggcgccgcucucgggggggcccaaguccuucuga-ucgaggccc--agcccgcggacggu   231
Fly           uaguaugagaaauuaacgauuuaaguccuucuuaaaugaggccauuuacccauagagggu   239
C.elegans     cagc--a-gugucguauguccgaaguccuuacga-uugaggcca-uaaaccagagagggu   225
Yeast         gggc--c-guuccuug--ucuauguuccuugg-------aacag-gacgucauagagggu   216
                *                      *****           *        *   ** ***

Human         gugaggccgguagcggccc-ccggcgcgc----cgggcccgggucuucccggagucgggu   286
Mouse         gugaggccgguagcggccc-ccggcgcgc----cgggcucgggucuucccggagucgggu   286
Chicken       gugaggccgguagcggccccccggcgcgc----cgggcccggggcuucucggagucgggu   287
Fly           gccaggcccguauaac------guuaaugauuacuagaugauguuuccaaagagucgugu   293
C.elegans     gcgagccccguucuggaua-gcggcacugu---ugguucgcuugcuccuuggagucgggu   281
Yeast         gagaaucccgug-uggcga-ggagugcgguucuuuguaaagugccuucgaagagucgagu   274
              *  *  ** **                                  * *   ****** **

Human         ugcuugggaaugcagcccaaagcggugguaaacuccaucuaaggcuaaauaccggcacg   346
Mouse         ugcuugggaaugcagcccaaagcggugguaaacuccaucuaaggcuaaauaccggcacg   346
Chicken       ugcuugggaaugcagcccaaagcggugguaaacuccaucuaaggcuaaauaccggcacg   347
Fly           ugcuugauagugcagcacuaagugggugguaaacuccaucuaaaacuaaauauaaccaug   353
C.elegans     ugcuugaaagugcagccuaaagugggugauaaacuucaucuaaggcuaaauaucgacucg   341
Yeast         uguuugggaaugcagcucuaagugggugguaaauuccaucuaaagcuaaauauuggcgag   334
              ** ***  * ******   *** ***** **** * *******  *******    *  *

Human         agaccgauagucaacaaguaccguaagggaaaguugaaaagaacuuugaagagagaguuc   406
Mouse         agaccgauagucaacaaguaccguaagggaaaguugaaaagaacuuugaagagagaguuc   406
Chicken       agaccgauagccaacaaguaccguaagggaaaguugaaaagaacuuugaagagagaguuc   407
Fly           agaccgauaguaaacaaguaccgugagggaaaguugaaaagaacucugaauagagaguua   413
C.elegans     auugcgauagcgaacaaguaccgugagggaaaguugcaaaggacuuugaagagagaguuc   401
Yeast         agaccgauagcgaacaaguacagugauggaaagaugaaaagaacuuugaaaagagaguga   394
              *   ******  ********* ** * ****** ** **** *** **** *******

Human         aagagggcgugaaaccguuaagagguaaacgggugggguccgcgcaguccgcccggagga   466
Mouse         aagagggcgugaaaccguuaagagguaaacgggugggguccgcgcaguccgcccggagga   466
Chicken       aagagggcgugaaaccguuaagagguaaacgggugggguccgcgcagucggcccggagga   467
Fly           aacaguacgugaaacugcuuagagguuaagcc-------------cgaugaaccugaaua   460
C.elegans     aagagaacgugaaaucgcuggaguggaaccggaga---------caguug------augu   446
Yeast         aaaaguacgugaaauuguugaaagggaaggg-------------cauuug------auca   435
              ** **  *******  * *     *  *                            *
```

```
Human       uucaacccggcggc--ggguccggccgugucggcgg-cccggcggaucuuucccgccccc    523
Mouse       uucaacccggcggcgcgcguccggccgugccgguggucccggcggaucuuucccgcuccc    526
Chicken     uucaacccggcgggccaaggucggccggcgcg--ggcgccgucggaucccc---g-----    517
Fly         ucc-------------g----------uuauggaa------------aauuca-------    478
C.elegans   ug---cuuggaga--caagcu---------uggug-------------acu---g-----    471
Yeast       -----------ga--ca-------------uggug-------------uuu---u-----    448
                                           *

Human       cguuccucccgaccccuccacccgcccucccuucccccgc--cgccccuccuccuccucc    581
Mouse       cguuccucccgaccccuccacccgcgcgucguuccccucuuccuccccgcguccggcgcu    586
Chicken     --------------ccuccgccuccccuccgucc--------cuccccu--------ucg    547
Fly         ------ucauua---------------------------------aa---auuguaauau    496
C.elegans   ------------------------------------------gucgcuuaguugugaucg    489
Yeast       ------------------------------------------gugcc----cucugcucc    462

Human       ccgga-gggggcgggcuccgg--cgggugcgg---gggugggcgggcggggccgggggug    635
Mouse       ccg----gcggcgggcgcggggggugguguggugguggcgcgcgggcggggccgggggug    642
Chicken     ccggggcggggcgggcccaggg------------ggg---------------------gc    574
Fly         uuaaau-------------------aauauuaugagaa--------------------ua    517
C.elegans   uugc---cggg-----------------------------------------------ug    499
Yeast       uugu---gggu-----------------------------------------------ag    472

Human       gggucggcgggggaccguccccccgaccggcgaccggccgccgccgggcgcauuuccaccg    695
Mouse       gggucggcgggggaccgccccccggccggcgaccggccgccgccgggcgcacuuccaccg    702
Chicken     gggcgggccggggaccgccgcccggccggcguccggcccccgucgggcgcauuuccuccg    634
Fly         ---------------------------------------------gugugcauuuuucca    533
C.elegans   ----------------------------------------------u--cguuuccuaug    511
Yeast       ----------------------------------------------gggaaucucgcauu    486
                                                             *

Human       cggcggugcgccgcgaccggcuccgggacggcugggaaggcccggcg--gggaagguggc    753
Mouse       uggcggugcgccgcgaccggcuccgggacggccgggaaggcccggug--gggaagguggc    760
Chicken     cggcggugcgccgcgaccggcuccgggacggcugggaagggcugccggcgggcagguggc    694
Fly         uaua--------aggacau------------------------------uguaa------    549
C.elegans   cuac-----g--ccgac-------------------------------------------    521
Yeast       ucac-----u--gggcca------------------------------------------    497
                          * *

Human       ucgggggggccccguccguccguccguccguccuccucccccgucuccgcccccggcccc    813
Mouse       ucggggggggcggcgcguc-------------------------uc--------------    781
Chicken     ccggcgccgcgcgagcggccg----ccgggugu-----------uauagccgccgggccc    739
Fly         -------------------------ucuauuagca---------uauacc----------    565
C.elegans   ------------------------------------------------------------    521
Yeast       ------------------------------------------------------------    497

Human       gcguccucccucggg--agggcgcgcgggucggggcggcggcggcgguggcggcggcggc    871
Mouse       -----------------agggcgcg-----------------------------------    789
Chicken     ggaucgucgccgaaucccggggccgagggagaggaccgccgccgcgccc--ucccccgga    797
Fly         aaauu-------------------------------------------------------    570
C.elegans   ------------------------------------------ggcguu------------    527
Yeast       gcauc-------------------------------------aguuuu------------    508

Human       ggcggcggcgggaccgaaaccccccccg-aguguuacagccccc---ccggcagcagcac    927
Mouse       --------ccgaac--caccucaccccg-aguguuacagcccu----ccggccgcagcuu    834
Chicken     gggggcggcccccc-ggagggcccccgcggccggaccggcgucgggccggccgcgccgc    856
Fly         --------uaucauaaaauauaacu-uauaguuuau------------------------    597
C.elegans   ---ggcugcucgu-----ucuagcccgacaguguugcccaucucg---------------    564
Yeast       ---gguggcagga-----uaaaucc-----------------------------------    525
                                   *
```

```
Human       ucgccgaaucccggggccgagggagcgag--acccgucgccgcgcucucc--ccccuccc   983
Mouse       ucgccgaaucccggggccgaggaagccagauacccgucgccgcgcucucc--cucuccc-   891
Chicken     gcgc--------gcguccgcgccgccgccguacgccgccgcucgcucucucuccguuccc   908
Fly         ---uccaauu---------------------aa-------auugcuu-----gcau----   617
C.elegans   ------------------------------------------------------------   564
Yeast       ------------------------------------------------------------   525

Human       ggc-----gcccacccccgcggggaaucccccgcgaggg-----gggucucccccgcggg   1033
Mouse       ccc-----guccgc----------------------------------cucccgggcgg-   911
Chicken     cgccccggguccgucccgggg----cgcgggggcggggggggucggguguccggcgcgcg   964
Fly         ---------uuuaa-----cacagaauaaau---------guuauuaauuugauaaagu-   653
C.elegans   ------------------caagagaag-----------------gugucuugcuggcgg-   588
Yeast       -------------------auaggaau-----------------guagcuugccucggu-   548
                                                             *       *

Human       ggcgcgccggcgucuccu--cgugggggggccg------------------ggccacccc   1073
Mouse       ---gcgug------------ggggugggggccg------------------ggccgcccc   938
Chicken     gcucggc--gcggcgccg--cgcguguggcgcgcgccuccagcccggcgcgggcgaggcc   1020
Fly         gcug----auagauuuauau-------gauu-------a---------------------   674
C.elegans   ---------uagug-------------gguucg---------------------------   599
Yeast       ---------aaguauuauagccugugggaauac---------------------------   572
                                       *

Human       ucc-------cacggcgcgaccgcucucccaccccuccuc--cccgcgcccccgcc----   1120
Mouse       ucc-------cacggcgcgaccgcucucccaccccccucc--gucgccuc---uc-----   981
Chicken     gcgggggcgccggggggggaaccuucccc--cuucuguucgggccgccuccguucccgcg   1078
Fly         ----------cagugcguuaauu-uuucggaauuau------------------------   699
C.elegans   ----------------------------------------------------uggcg---   604
Yeast       ----------------------------------------------------ugcca---   577

Human       -------------ccggcgacggggggggu---gccgcgcgcgggucggggggcggggcg   1164
Mouse       -------------ucgg----------------------ggcccgguggggggcggggcg   1006
Chicken     ggggcggcccguucgggggacgggcccgccggcccccggcgccgcuguccgaccggggcg   1138
Fly         ---------------------------------------------------auaauggca   708
C.elegans   ---------------------------------------------------gcuagcguu   613
Yeast       ---------------------------------------------------gcugggacu   586

Human       gacuguccccagugcgccccgggcg-ggucgcgccgucgggcccgggggagguucucucg   1223
Mouse       gacuguccccagugcgccccgggcgucgucgcgccgucggucccgggggggaccgu--cg   1064
Chicken     gacugcgcucagugcgccccgaccg-cgcggcgccgccgggccgg----------gcucg   1187
Fly         uaauua--ucauugau----------uuuug------------------------uguuu   732
C.elegans   uaguuacgcuagu-------------gugugugacgucgg---------------ugu--   643
Yeast       ga----------g-------------gacugcgacgu-----------------------   600
             *                            *

Human       gggccacgcgcgcgucccccgaagagggggacggcggagcgagcgcacggggucggcggc   1283
Mouse       --g----ucacgcgucucccgacgaag-----------ccgagcgcacggggucggcggc   1107
Chicken     g-gccacgc-----------------------------cagggcgcccgggguccgcggc   1217
Fly         auuauaugcacuuguaugauuaacaaugcga---------------------------aa   765
C.elegans   --gaaagucg-----------acga-----------------------------------   655
Yeast       ----aaguca-----------aggau------------gcuggc----------------   617
        *

Human       gacgucggcuacccacccgacccgucuugaaacacggaccaaggagucuaacacgugcgc   1343
Mouse       gaugucggcuacccacccgacccgucuugaaacacggaccaaggagucuaacgcgugcgc   1167
Chicken     gacgucggcuacccacccgacccgucuuguaacacggaccaaggagucuagcacgcgcgc   1277
Fly         gauucaggauaccuucgggacccgucuugaaacacggaccaaggagucuaacauaugugc   825
C.elegans   -----------cguuuccgacccgucuugaaacacggauugcggagugcuugucuacugc   704
Yeast       -auaaugg--uuauaugccgcccgucuugaaacacggaccaaggagucuaacgucuaugc   674
                                ********* ********    *****           **
```

Human gagucgggggcucgcacgaaagccgccguggcgcaaugaaggugaaggccggcgcgcucg 1403
Mouse gagucaggggcucguccgaaagccgccguggcgcaaugaaggugaagggccccgcccg-g 1226
Chicken gagucggcggcucgcgcgaaagcc--cgcggcgcaaugaaggugagggccggcgcgc--g 1333
Fly aaguuauugggau-----auaaaccuaauagcguaauuaacuugacuaauaaugggauua 880
C.elegans gagucaaagggugu----uaaaaccuugcggcgaaaugaaaguaaaggucagucucg-aa 759
Yeast gaguguuugggugu----aaaacccaua-cgcguaaugaaagugaacguagguuggg-gc 728
 ***    **          *  *      *** *** **  * *

Human ccggccgaggu-----------------gggaucccgaggccuc---uccaguccgccga 1443
Mouse gggcccgaggu-----------------gggaucccgaggccuc---uccaguccgccga 1266
Chicken ccggcugaggu-----------------gggaucccggggcggca-ggccggaaggcccc 1375
Fly guuuuuuagcuauuuauagcuaauuaacacaaucccggggcguucuauauaguu------ 934
C.elegans uuggccgacgu-----------------gggaucuguguucuucg----------gagug 792
Yeast cucgc-----------------------aag----------------------------- 736

Human gggcgcaccaccggcccgucucgcccgccgcgccggggagg-----uggagcacgagcgc 1498
Mouse gggcgcaccaccggcccgucucgcccgccgcgccggggagg-----uggagcacgagcgu 1321
Chicken gggcgcaccaccggcccgucucgcccgccucgccggggagg-----uggagcaugagcgc 1430
Fly --auguauaauguau--auuuauauuauuuaugccucuaacuggaacguaccuugagcau 990
C.elegans cagcgcaccacggcccugugcgugucacuugug--acug----------ugcagagguug 840
Yeast aggugcacaaucgaccgauccugaugucuucgg--auggau-----uugaguaagagcau 789
    * *  *        *                                     *

Human acguguua-----ggacccgaaagauggugaacuaugccugggcagggcgaagccagagg 1553
Mouse acgcguua-----ggacccgaaagauggugaacuaugccugggcagggcgaagccagagg 1376
Chicken gcgugcua-----ggacccgaaagauggugaacuaugccugggcagggcgaagccagagg 1485
Fly auaugcug-----ugacccgaaagauggugaacuauacuugaucagguugaagucagggg 1045
C.elegans agcaguuggcaaacgacccgaaagauggugaacuaugccugagcaggaugaagccagagg 900
Yeast agcuguug-----ggacccgaaagauggugaacuaugccugaauagggugaagccagagg 844
    * *       ********************** * **   ***  **** *** **

Human aaacucugguggagguccguagcgguccugacgugcaaaucggucguccgaccuggguau 1613
Mouse aaacucugguggagguccguagcgguccugacgugcaaaucggucguccgaccuggguau 1436
Chicken aaacucugguggagguccguagcgguccugacgugcaaaucggucguccgacccggguau 1545
Fly aaacccugauggaagaccgaaacaguucugacgugcaaaucgauugucagaauugaguau 1105
C.elegans aaacucugguggaaguccguaucgguucugacgugcaaaucgaucgauagacuuggguau 960
Yeast aaacucugguggaggcucguagcgguucugacgugcaaaucgaucgucgaauuuggguau 904
**** *** **** *  ** * * ** *************** * *    *   * ****

Human aggggcgaaagacuaaucgaaccaucuaguagcugguucccuccgaaguuucccucagga 1673
Mouse aggggcgaaagacuaaucgaaccaucuaguagcugguucccuccgaaguuucccucagga 1496
Chicken aggggcgaaagacuaaucgaaccaucuaguagcugguucccuccgaaguuucccucagga 1605
Fly aggggcgaaagaccaaucgaaccaucuaguagcugguuccuuccgaaguuucccucagga 1165
C.elegans aggggcgaaagacuaaucgaaccaucuaguagcugguuccuuccgaaguuucccucagga 1020
Yeast aggggcgaaagacuaaucgaaccaucuaguagcugguuccugccgaaguuucccucagga 964
************* **************************  ******************

Human uagcuggcgcucucgcagacccgacgcacccccgccacgcaguuuuauccgguaaagcga 1733
Mouse uagcuggcgcucucgcu--cccgac---------guacgcaguuuuauccgguaaagcga 1545
Chicken uagcuggcgcucggggcgg---------------cggugcaguuuuacccgguaaagcga 1650
Fly uagcuggugcauuuuaauauuaua----------uaaaauaaucuuaucugguaaagcga 1215
C.elegans uagcuggaucuc-----------------------ag-gcaguuauauucgguaaagcua 1056
Yeast uagcagaagcuc-----------------------guaucaguuuuaugagguaaagcga 1001
**** *   *                              * *  **   ******** *

Human augauuagaggucuuggggccgaaacgaucucaaccuauucucaaacuuuaaaugggaa 1793
Mouse augauuagaggucuuggggccgaaacgaucucaaccuauucucaaacuuuaaaugggaa 1605
Chicken augauuagaggucuuggggccgaaacgaucucaaccuauucucaaacuuucaaugggaa 1710
Fly augauuagaggccuuagggucgaaacgaucuuaaccuauucucaaacuuuaaaugggaa 1275
C.elegans augauuagaggccuuggggacguaauguccucaaccuauucucaaacuuucaauggauau 1116
Yeast augauuagagguuccgggucgaaaugaccuugaccuauucucaaacuuuaaauauguaa 1061
***********     *** ** ** *  **  ****************  ***   **

```
Human       gaagcccggcucgcuggcgugg-agccggg-cguggaaug-cgagugccuagugggccac    1850
Mouse       gaagcccggcucgcuggcgugg-agccggg-cguggaaug-cgagugccuagugggccac    1662
Chicken     gacgcccggcucgcuggcgugg-agccgggccguggaaug-cgagcgcucagugggccac    1768
Fly         gaaccuuaacuuucuugauaugaaguucaagguuaugauauaaugugcccagugggccac    1335
C.elegans   gaaguugcaguuucuuuaguga-acugu-caacgugaaug-cgagguccaagugggccau    1173
Yeast       gaaguccuuguuacuuaauuga-acguggacauuugaaugaagagcuuuuagugggccau    1120
            **        *  **        *             **     *     *********

Human       uuuugguaagcagaacuggcgcugcgggaugaaccgaacgccggguuaaggcgcccgaug    1910
Mouse       uuuugguaagcagaacuggcgcugcgggaugaaccgaacgccggguuaaggcgcccgaug    1722
Chicken     uuuugguaagcagaacuggcgcugcgggaugaaccgaacgccggguuaaggcgcccgaug    1828
Fly         uuuugguaagcagaacuggcgcugugggaugaaccaaacguaauguuacggugcccaaau    1395
C.elegans   uuuugguaagcagaacuggcgcugugggaugaaccaaacguggaguuaaggugccuaacu    1233
Yeast       uuuugguaagcagaacuggcgaugcgggaugaaccgaacguagaguuaaggugccggaau    1180
            ********************* ** ********** ****    **** ** ***  *

Human       ccgacgcucau-cagaccccagaaaagguguugguugauauagacagcaggacgguggcc    1969
Mouse       ccgacgcucau-cagaccccagaaaagguguugguugauauagacagcaggacgguggcc    1781
Chicken     ccgacgcucau-cagagcccagaaaagguguugguugaucuagacagcaggacgguggcc    1887
Fly         uaacaacucaugcagauaccaugaaaggcguugguugcuuaaaacagcaggacggugauc    1455
C.elegans   -ucucgcucau-gagaccccauaaaagguguugguugauauugacagcaggacgguggcc    1291
Yeast       -acacgcucau-cagacaccacaaaagguguuaguucaucuagacagccggacgguggcc    1238
                  *****  ***  ***  ***** *** ***  *    ***** ********  *

Human       auggaagucggaauccgcuaaggaguguguaacaacucaccugccgaaucaacuagcccu    2029
Mouse       auggaagucggaauccgcuaaggaguguguaacaacucaccugccgaaucaacuagcccu    1841
Chicken     auggaagucggaacccgcuaaggaguguguaacaacucaccugccgaaucaacuagcccu    1947
Fly         auggaagucgaaauccgcuaaggaguguguaacaacucaccugccgaagcaacuagcccu    1515
C.elegans   auggaagucgguauccgcuaaggaguguguaacaacucaccugccgaaucaacuagcccu    1351
Yeast       auggaagucggaauccgcuaaggaguguguaacaacucaccggccgaaugaacuagcccu    1298
            **********  * *************************** ******  **********

Human       gaaaauggauggcgcuggagcgucgggcccauacccggccgucgccggcagucgagagug    2089
Mouse       gaaaauggauggcgcuggagcgucgggcccauacccggccgucgccggcagucggaacgg    1901
Chicken     gaaaauggauggcgcuggagcgucgggcccauacccggccgucgccggcggugcgga---    2004
Fly         uaaaauggauggcgcuuaaguuguauaccuauacauuaccgcuaaaguagauga------    1569
C.elegans   gaaaauggauggcgcuuaagcgagagaccuauacuccgccguugcgacaugu--------    1403
Yeast       gaaaauggauggcgcucaagcguguuaccuauacucuaccgucagggu------------    1346
             ***************  **       ** ****    ***

Human       gacgggagcggcgggggcggcgcgcgcgcgcgcguguggugugcgucggagggcggcg    2149
Mouse       aacgggacgggagcggc-------cgcgggugcgcgucucuc-ggggucggggg------    1947
Chicken     ------------------------------------------------------------    2004
Fly         ------------------------------------uuuaua-uuacuug----------    1582
C.elegans   --------------------------------------------gcguug----------    1409
Yeast       ------------------------------------------------ug----------    1348

Human       gcggcggcggcggcggggguguggggguccuucccccgcccccccccccccacgccuccucc    2209
Mouse       -------------ugcguggcgggggcccgucccccgccuccccuccgcgcgccggguu-    1993
Chicken     ------------------------------------------------------------    2004
Fly         ------------------------------------------------------------    1582
C.elegans   ------------------------------------------------------------    1409
Yeast       ------------------------------------------------------------    1348

Human       ccuccucccgcccacgccccgcuccucgcccccggagccccgcggacgcuacgccgcgac    2269
Mouse       -------ucgccccg-----------cg-gcgucgggccccgcggacgcuacgccgcgac    2035
Chicken     --------------------------------------gccgcgggggcuacgccgcgac    2026
Fly         ----------------------------ugau------a--uaaauuuugaaacuuuagu    1606
C.elegans   -----------------------------------------ucuagcgccaggucguaac    1428
Yeast       -----------------------------------------a----uaugaugcccugac    1363
                                                              *
```

```
Human      gaguaggagggccgcugcggugagccuugaagccu-agggcgcgggcccggguggagccg   2328
Mouse      gaguaggagggccgcugcggugagccuugaagccu-agggcgcgggcccggguggagccg   2094
Chicken    gaguaggagggccgcugcggugcgccuggaagccu-ggggcgcgggcccggguggagccg   2085
Fly        gaguaggaagg-uacaaugguaugcguagaagugu-uuggcguaagccugcauggagcug   1664
C.elegans  gaguaggaaggucguggcgguu-gcguugaaggcuaugagcguaggcucggcuggagcuu   1487
Yeast      gaguaggcagg-cguggaggucagugacgaagccu-agaccguaaggucgggucgaacgg   1421
           *******  **       ***  *    ****  *     **   *   *  * ** *

Human      ccgcaggugcagaucuuggugguaguagcaaauauucaaacgagaacuuugaaggccgaa   2388
Mouse      ccgcaggugcagaucuuggugguaguagcaaauauucaaacgagaacuuugaaggccgaa   2154
Chicken    ccgcaggugcagaucuuggugguaguagcaacuauucaaacgagagcuuugaaggccgaa   2145
Fly        ccauugguacagaucuuggugguaguagcaaauaaucgaaugagaccuuggaggacugaa   1724
C.elegans  ccgucagugcagaucguaaugguaguagcaaauauucaaguucgauccuugaagacugaa   1547
Yeast      ccucuagugcagaucuuggugguaguagcaaauauucaaaugagaacuuugaagacugaa   1481
           **    ** ****** *  ************ ** ** *    ** * * ** * * ***

Human      guggagaaggguuccaugugaacagcaguugaacaugggucagucgguccugagagaugg   2448
Mouse      guggagaaggguuccaugugaacagcaguugaacaugggucagucgguccugagagaugg   2214
Chicken    guggagcaggguuccaugugaacagcaguugaacaugggucagucgguccuaagcgauag   2205
Fly        guggagaaggguuucgugugaacagugguugaucacgaguuagucgguccuaaguucaag   1784
C.elegans  guggagaaggguuccacgugaacaguaguuggaugugggucagucgauccuaagguacug   1607
Yeast      gugggaaagguuccacgucaacagcaguuggacgugggguuagucgauccuaagagaugg   1541
           **** * * **** *  ** *****  ****     * ** ***** **** **     *

Human      gcgagcgccguu----ccgaagggacg-ggcgauggccucc-------------------   2484
Mouse      gcgagugccguu----ccgaagggacg-ggcgauggccucc-------------------   2250
Chicken    gcgagcgccguu----ccgaagggacg-ggcgauggccucc-------------------   2241
Fly        gcgaaagccgaaaauuuucaaguaaaacaaaaauggcuaacuauauaaacaaagcgaauu   1844
C.elegans  gcgaacgccuug----uaucaucggug-gcgaaaagcuugc-------------------   1643
Yeast      ggaagcuccguu------u-----------caaaggccuga-------------------   1565
           *  *   **                       *  **

Human      --------guugc--ccucggccgaucgaaagggagucggguucagauccccgaauccgg   2534
Mouse      --------guugc--ccucggccgaucgaaagggagucggguucagauccucgaauccgg   2300
Chicken    --------guugc--ccucagccgaucgaaagggagucggguucagauccccgaauccgg   2291
Fly        auaauacacuugaaua--auuuugaacgaaagggaauacgguuccaauuccguaaccugu   1902
C.elegans  --------uuuuagucc-ccgcuugucgaaagggaauaggguuaauauucccuaacugag   1694
Yeast      --------uuuuaugcaggccaccaucgaaagggaauccgguuaagauuccggaaccugg   1617
                    **               ********* *  ****   ** *   **

Human      agug-------------gcggagaugggcgccgcgaggcguccagugcgguaacgcgacc   2581
Mouse      agug-------------gcggagaugggcgccgcgaggcguccagugcgguaacgcgacg   2347
Chicken    agcg-------------gcggagacgggcgccgcgaggcgcccagugcgguaacgcaagc   2338
Fly        ugaguauccguuuguuauuaaauaugggccucgu-----gcucauccuggcaacaggaac   1957
C.elegans  au---------------gcaaagauuguguucuucggagcacaagcgcgguaacgcauuc   1739
Yeast      au---------------au-------ggauucuuc-----------acgguaacguaacu   1644
                                     *    *                **  ***

Human      gaucccggagaagccggcgggagccccggggagaguucucuuuucuuugugaagggcagg   2641
Mouse      gaucccggagaagccggcgggagccccggggagaguucucuuuucuuugugaagggcagg   2407
Chicken    gaucccggagaagccggcgggagccccggggagaguucucuuuucuuugugaagggccgg   2398
Fly        gaccauaaagaagccgucgagagauaucggaagaguuuucuuuucuguuuuauagccgua   2017
C.elegans  gaacuugguua-gucgcucaaagaccgagcuagaguuuucuucucua-guuaaggaacgg   1797
Yeast      gaauguggagacgucggcgcgagcccugggaggaguuaucuuuucuu-cuuaacagcuua   1703
           **        * * **     **     *   ***** **** ***   * *

Human      gcgcccuggaaugguucgccccgagagaggggcccgugccuuggaaagcgucgcgguuc   2701
Mouse      gcgcccuggaaugguucgccccgagagaggggcccgugccuuggaaagcgucgcgguuc   2467
Chicken    gcgcccuggaacgguucgccccgagagaggggcccgcgccuuggaaagcgucgcgguuc   2458
Fly        cuaccauggaagucuuucgcagagagagauaugguagaugggcuagaagagcaugacauaua   2077
C.elegans  acucccuggaauugguucagccagagaugggacguuguuuccgaaaagcaccgcgguuu   1857
Yeast      ucaccccggaauugguuuauccggagaugggucuuauggcuggaagaggccagcaccuu   1763
              **  ****    **      ****   **           * * **     *   *
```

```
Human       cggcggcguccggugagcucucgcuggcccuugaaaauccgggggagagggug-uaaauc     2760
Mouse       cggcggcguccggugagcucucgcuggcccuugaaaauccgggggagagggug-uaaauc     2526
Chicken     cggcggcguccggugagcucucgcuggcccgugaaaauccgggggagagggug-uaaauc     2517
Fly         cuguuguguc-gauauuuucuccucggaccuugaaaauuuaugguggggacacgcaaacu     2136
C.elegans   cuguggugucucgugcucuuugaacggcccuuaaaacaccaagggag--gcuauuaauuu     1915
Yeast       -ugcuggcuccggugcgcuugugacggcccgugaaaauccacaggaa--ggaa-uaguuu     1819
              *  *  **   *    *      ** ** * ***       *           *

Human       ucgcgccgggccguacccauauccgcagcaggucuccaaggugaacagccucuggcaugu     2820
Mouse       ucgcgccgggccguacccauauccgcagcaggucuccaaggugaacagccucuggcaugu     2586
Chicken     ucgcgccgggccguacccauauccgcagcaggucuccaaggugaacagccucuggcaugu     2577
Fly         ucucaacaggccguaccaauauccgcagcuggucuccaaggugaagagucucuagu-cga     2195
C.elegans   gcacu--caaucguaccgauauccgcauuaggucuccaaggugaacagccucuagu-cga     1972
Yeast       ucaugccaggucguacugauaaccgcagcaggucuccaaggugaacagccucuagu-uga     1878
             *         *****  *** *****   *************** ** **** *   *

Human       uggaacaauguagguaagggaagucggcaagccggauccguaacuucgggauaaggauug     2880
Mouse       uggaacaauguagguaagggaagucggcaagccggauccguaacuucgggauaaggauug     2646
Chicken     uggaccaauguagguaagggaagucggcaagccggauccguaacuucgggauaaggauug     2637
Fly         uagaauaauguagguaagggaagucggcaaauuagauccguaacuucgggauaaggauug     2255
C.elegans   uagaauaauguagguaagggaagucggcaaacuagauccguaacuucgggaaaaggauug     2032
Yeast       uagaauaauguagauaagggaagucggcaaaauagauccguaacuucgggauaaggauug     1938
            * **  ******* ****************    ****************  ********

Human       gcucuaagggcugggucggucgggcuggggcgcgaagcggggcugggcgcgcgccgcggc     2940
Mouse       gcucuaagggcugggucggucgggcuggggcgcgaagcggggcugggcgcgcgccgcggc     2706
Chicken     gcucuaagggcugggucggucgggcuggggcgcgaagcggggcugggcgcgcgccgcggc     2697
Fly         gcucugaagauugagauagucgggcuugauugggaaacaau-------------------     2296
C.elegans   gcuccagugguuggaacgguuggccaguug------------------------------     2062
Yeast       gcucuaagggucgguagugagggcc-uug-------------------------------     1967
            ****    *   *        ** *

Human       uggacgaggcgccgccgccccc-cccacgcccggggcacccc-ccuc----gcggcccuc     2994
Mouse       uggacgaggcgccgccgcccucucccacguccggggagaccccccguccuuuccgcccgg     2766
Chicken     uggacgaggcgccgcccgcccccgc------------------ccccccuuuc-------     2732
Fly         -----a-------------------------------------acaugguuuaugugcu-     2313
C.elegans   --guugaugcuuguccggc--------------------------gcaguucu-g-----     2088
Yeast       --gu-cagacgcagcgggc--------------------------gugcuugugg-----     1993

Human       ccccgcccc-------accccgcgcgcgccgcucgcucccuccccaccccgcgcccucuc     3047
Mouse       gcccgcccuccccucuuccccgcggggcc---------ccgucgucccccgcgucgucgc     2817
Chicken     ------------------------------------------------------------     2732
Fly         ---------------------------------cguucug--gguaaauaga---guuuc     2335
C.elegans   ------------------------------------------------------------     2088
Yeast       ------------------------------------------------------------     1993

Human       ucucucucucucccccgcucccc----guccuccccccuccccgggggagcgccgcgugg     3103
Mouse       caccucucuuccc-ccc----------cuccuucuucccgucggggggcgg-----gucg     2861
Chicken     -------------cccgcucccg----cucg---------ccg----gggcgccgggggg     2762
Fly         u------------agcauuuauguuaguuacuuguuc-----------------------     2360
C.elegans   -------------ucugcuu--g----auacu--uu----------------------cg     2105
Yeast       -------------acugcuu--g----guggggcuu----------------------gc     2012
                                        *

Human       gggcggcggcggggggagaagggucggg-gcggcaggggccggcggcggcccgccgcggg     3162
Mouse       gg-ggucggcgcgcggcgcgggcuccggggcggcgg-gucc--------aaccccgcggg     2911
Chicken     gggggu------------------------------------------------------     2768
Fly         ------------------------------------------------------------     2360
C.elegans   gguugaug---------------------gc----------------g------------     2116
Yeast       ucugcuag---------------------gc----------------g------------     2023
```

```
Human       g-ccccggcggcgggggcac----gguc-ccc---ugcgaggggggcccgggcacccgg-    3212
Mouse       gguuccggagcgggaggaaccagcgggcccccgguggggcggggggcccggacacucggg    2971
Chicken     ----------------------------------------------------ca----gc    2772
Fly         -----------------------------ccc-----------------ggauaguuuag    2374
C.elegans   -----------------gac--------------------------------uagug---    2124
Yeast       -----------------gac--------------------------------uacuu---    2031
                                                                 *

Human       ggggccggcggcggcggcgacucuggacgcgagccgggcccuucccguggaucgccccag    3272
Mouse       ggggccggcggcggcggcgacucuggacgcgagccgggcccuucccguggaucgccucag    3031
Chicken     gggcggcgcggcggcggcgacucuggacgcgcgccgggcccuucccguggaucgccccag    2832
Fly         u--------uacguagccaauugugg-----------aacuuucuugcuaaaauu-----    2410
C.elegans   -------------------auu--------------gugcuugcuugcggacgcuu----    2147
Yeast       -------------------gc---------------gugccuuguuguagacggcc----    2053
                                                   * *    *   *

Human       cugcggcgggcgucgcggccgcccccggggagcccggcgggcgccggcgcgccccccccc    3332
Mouse       cugcggcgggcgucgcggccgcucccggggagcccggcgggugccggcgcggguccccuc    3091
Chicken     cugcggcgggcgccgcucgcccccucc-----------uugccccuccgc----------    2872
Fly         ------------------------------------------------------------    2410
C.elegans   ------------------------------------------------------------    2147
Yeast       ------------------------------------------------------------    2053

Human       acccccaccccacgucucgucgcgcgcguccgcugggggcggggagcggucgggcggc      3392
Mouse       cccgcgggg--------ccu----------------------------------------    3103
Chicken     --------ccccccgcucccg--gcgccccuccc--------------------------    2896
Fly         -----------------uuua-aga----a------------------------------    2418
C.elegans   -----------------ucuggugu-------------gugc------------------    2159
Yeast       -----------------uu-----------------------------------------    2055

Human       ggcggucggcgggcggcggggcggggcgguucguccccccgcccuacccccccggccccg    3452
Mouse       ---------------------------------------cgcuccacccc----------    3114
Chicken     ----gucggccg------------------ucgucc---cggccg---ccc-----cccg    2923
Fly         ------------------------------uacuauuuggguuaaa--------------    2434
C.elegans   ------------------------------uuggaccucgguucuaguauc---------    2180
Yeast       -------------------------------gguaggucucuuguagaccg---------    2075

Human       uccgccccccguuccccccuccuc-------cucggcgcgcggcggcggcggcggc-gg-    3503
Mouse       --------cccaucgccucu--cc-------cgaggugcguggcgggggcgggcgggcgu    3157
Chicken     uc------ccgagcg-cccuccucgcgagggcgcgaggggcggcggcggccgcgggc    2976
Fly         --------ccaauuaguucuuauu------------------------------------    2450
C.elegans   --------cugauc----------------------------------------------    2186
Yeast       --------ucgcuu----------------------------------------------    2081

Human       -cggcggg---cggcggaggggccgcgggccggu-ccccccgccggguccgccccc--g    3556
Mouse       cccgcgcgugugggggaaccuccg--cgucgguguuccccgccggguccgccccc--c    3213
Chicken     gcggcggcggcggg-ggg----gggcccgccgguggcgccgggcgg-ggcggucccgggc    3030
Fly         ------------------------------------------------------------    2450
C.elegans   ------------------------------------------------------------    2186
Yeast       ------------------------------------------------------------    2081

Human       gggccgcggu--uccgcgcggcgccucgccucggccggcgccuagcagccgacuuagaac    3614
Mouse       gggccgcgguuuuccgcgcggcgccucgccucggccggcgccuagcagccgacuuagaac    3273
Chicken     gggggggggucuccgggccggcgccccgccucggccggcgccuagcagccggcuuagaac    3090
Fly         -------------aa----------uuauaacgauuaucaauuaacaaucaauucagaac    2487
C.elegans   ----------------------------------gcucaucuaaacaaccguacuggaac    2212
Yeast       ----------------------------------gcuacaauuaacgaucaacuuagaac    2107
                                                       * *   *      ****
```

```
Human       uggugcggaccaggggaauccgacuguuuaauuaaaacaaagcaucgcgaaggcccgcgg   3674
Mouse       uggugcggaccaggggaauccgacuguuuaauuaaaacaaagcaucgcgaaggcccgcgg   3333
Chicken     uggugcggaccaggggaauccgacuguuuaauuaaaacaaagcaucgcgaaggcccgcgg   3150
Fly         uggcacggacuuggggaauccgacugucuaauuaaaacaaagcauugugauggcccuag-   2546
C.elegans   cgguacggacucagggaauccgacugucuaauuaaaacagaggugacagauggucc uug-  2271
Yeast       ugguacggacaaggggaaucugacugucuaauuaaaacauagcauugcgauggucagaaa   2167
             **  *****   ******* ****** *********** **      ** ** *

Human       cggguguugacgcgaugugauuucugcccagugcucugaaugucaaagugaagaaauuca   3734
Mouse       cggguguugacgcgaugugauuucugcccagugcucugaaugucaaagugaagaaauuca   3393
Chicken     cggguguugacgcgaugugauuucugcccagugcucugaaugucaaagugaagaaauuca   3210
Fly         cggguguugacacaaugugauuucugcccagugcucugaaugucaaagugaagaaauuca   2606
C.elegans   cggacguugacugucacugauuucugcccagugcucugaauguuaaaucguaguaauucg   2331
Yeast       gugauguugacgcaaugugauuucugcccagugcucugaaugucaaagugaagaaauuca   2227
              *  ******      ************************** ***  * ** *****

Human       augaagcgcggguaaacggcgggaguaacuaugacucucuuaagguagccaaaugccucg   3794
Mouse       augaagcgcggguaaacggcgggaguaacuaugacucucuuaagguagccaaaugccucg   3453
Chicken     augaagcgcggguaaacggcgggaguaacuaugacucucuuaagguagccaaaugccucg   3270
Fly         aguaagcgcgggucaacggcgggaguaacuaugacucucuuaagguagccaaaugccucg   2666
C.elegans   aguaagcgcggguaaacggcgggaguaacuaugacucucuuaagguagccaaaugccucg   2391
Yeast       accaagcgcggguaaacggcgggaguaacuaugacucucuuaagguagccaaaugccucg   2287
            *  ********** **********************************************

Human       ucaucuaauuagugacgcgcaugaauggaugaacgagauucccacugucccuaccuacua   3854
Mouse       ucaucuaauuagugacgcgcaugaauggaugaacgagauucccacugucccuaccuacua   3513
Chicken     ucaucuaauuagugacgcgcaugaauggaugaacgagauucccacugucccuaccuacuc   3330
Fly         ucaucuaauuagugacgcgcaugaauggauuaacgagauuccuacugucccuaucuacua   2726
C.elegans   ucauuuaauuguugacgcgcaugaauggauuaacgagauuccuacugucccuaacuacuu   2451
Yeast       ucaucuaauuagugacgcgcaugaauggauuaacgagauucccacugucccuaucuacua   2347
            **** *****  ****************** *********** ********** *****

Human       uccagcgaaaccacagccaagggaacgggcuuggcggaaucagcggggaaagaagacccu   3914
Mouse       uccagcgaaaccacagccaagggaacgggcuuggcggaaucagcggggaaagaagacccu   3573
Chicken     uccagcgaaaccacagccaagggaacgggcuuggcggaaucagcggggaaagaagacccu   3390
Fly         ucuagcgaaaccacagccaagggaacgggcuuggaauaauuagcggggaaagaagacccu   2786
C.elegans   ucuagcgaaaccacagccaagggaacgggcuuggcaaaaauagcggggaaagaagacccu   2511
Yeast       ucuagcgaaaccacagccaagggaacgggcuuggcagaaucagcggggaaagaagacccu   2407
            ** *******************************   **  *******************

Human       guugagcuugacucuagucuggcacggugaagagacaugagagguguagaauaagugggа   3974
Mouse       guugagcuugacucuagucuggcacggugaagagacaugagagguguagaauaaguggga   3633
Chicken     guugagcuugacucuagucuggcgcugugaagagacaugagagguguagaauaaguggga   3450
Fly         uuugagcuugacucuaaucuggcaguguaaggagacauaagagguguagaauaaguggga   2846
C.elegans   guugagcuugacucuaguuugacauugugaagagucaugagagguguagcauagguggga   2571
Yeast       guugagcuugacucuaguuugacauugugaagagacauagaggguguagaauaaguggga   2467
             *************** * ** *   ** * *** ***    ******* *** ******

Human       ggccccggcgcccccccggugucccgcgaggggcccggggcggggu cgccggcccug   4034
Mouse       ggccccggcgcccggcc-ccguccucgcgu-----cggggucggggcacgccggccucg   3687
Chicken     ggccccgcggucg-------------cgcga----cccgcgccgcg------------gc   3481
Fly         gauauuagacc--------------ucg------------g----------------uuu   2864
C.elegans   gucuuc------------------------------------------------------   2577
Yeast       gcuu--------------------------------------------------------   2471
            *

Human       cgggccgccggugaaauaccacuacucugaucguuuuuucacugacccggugaggcgggg   4094
Mouse       cgggccgccggugaaauaccacuacucucaucguuuuuucacugacccggugaggcgggg   3747
Chicken     ccggccgccggugaaauaccacuacucugaucguuuuuucacuuacccggugaggcgggg   3541
Fly         gguaucgucaaugaaauaccacuacucuuauuguuuccuuacuuacuugauuaaauggaa   2924
C.elegans   --ggacgacagugaaauaccaccacuuucaucgacucuuuacuuauucgguuaaaagaga   2635
Yeast       --cggcgccagugaaauaccacuaccuuuauaguuucuuuacuuauucaaugaagcggag   2529
                 ** *  *********** **  * ** *  *  * *** *     * *   *
```

```
Human       gggcgagc---cccgagggg------------------------------------cu--   4113
Mouse       gggcgagc---cccgagggg------------------------------------cu--   3766
Chicken     gggcgagc---cccgagggg------------------------------------cu--   3560
Fly         cguguaucauuuccuagccauuauacggauauauuuauuauaucuuaugguauuggguuu   2984
C.elegans   auuggcuu---cacg---------------------------------------------   2647
Yeast       cuggaauu---cauu---------------------------------------------   2541

Human       -----cucgcuucuggcgccaagcgcccggcc----------------------------   4140
Mouse       -----cucgcuucuggcgccaagcguccgucc----------------------------   3793
Chicken     -----cucgcuucuggcgccaagcgcccggcg----------------------------   3587
Fly         ugaugcaagcuucuugaucaaaguaucacgaguuuguuauauaaucgcaaacaaauucuu   3044
C.elegans   ---------gccuuuuuucgaagcauuaagcg----------------------------   2670
Yeast       ---------uuccacguuc-uagcauucaag-----------------------------   2562
                              *  **

Human       ----------gcgcg--------------------------------------cc-ggcc   4151
Mouse       ---------cgcgcgu-------------------------------------gc-gggc   3806
Chicken     ---------cgc---------------------------------------------gcc   3593
Fly         uaauaaaacgaugcauuuauguauuuuugauuugaaaauugguauaacuccaauuacuc   3104
C.elegans   ---------gagccau-------------------------------------uuuaugg   2684
Yeast       ---------gucccau-------------------------------------ucg----   2572

Human       gggcgcgacccgcuccggggacagugccagguggggaguuugacuggggcgguacaccug   4211
Mouse       gggcgcgacccgcuccggggacagugccagguggggaguuugacuggggcgguacaccug   3866
Chicken     gggcgcgacccgcuccggggacagcgucagguggggaguuugacuggggcgguacaccug   3653
Fly         agguaugauccaauucaaggacauugccagguagggaguuugacuggggcgguacaucuc   3164
C.elegans   caccgugacucuccucgaagacagugucaagcggggaguuugacuggggcgguacaucua   2744
Yeast       -gggcugauccggguugaagacauugucagguggggaguuggcuggggcggcacaucug   2631
                  **  *        ****  * ** *  ********* ********* *** **

Human       ucaaacgguaacgcagguguccuaaggcgagcucagggaggacagaaaccucccguggag   4271
Mouse       ucaaacgguaacgcagguguccuaaggcgagcucagggaggacagaaaccucccguggag   3926
Chicken     ucaaagcguaacgcagguguccuaaggcgagcucagggaggccagaaaccucccguggag   3713
Fly         ucaaauaauaacggagguguccca aggccagcucagugcggacagaaaccacacauagag   3224
C.elegans   ucaaaucguaacguagguguccuaaggcgagcucagagaggacggaaaccucucguagag   2804
Yeast       uuaaacgauaacgcagauguccuaaggggggcucauggagaacagaaaucuccaguagaa   2691
            * ***   ***** ** ***** ****   *****  * *  * **** * *   * **

Human       cagaagggcaaaagcucgcuugaucuugauuuucaguacgaauacagaccgugaaagcgg   4331
Mouse       cagaagggcaaaagcucgcuugaucuugauuuucaguacgaauacagaccgugaaagcgg   3986
Chicken     cagaagggcaaaagcucgcuugaucuugauuuucaguacgaauacagaccgugaaagcgg   3773
Fly         caaaagggcaaaugcugacuugaucucgguguucaguacacacagggacagcaaaagcuc   3284
C.elegans   caaaagggcaaaagcuugcuugaucuugacuuucaguacgaguacagaccgcgaaagcgu   2864
Yeast       caaaaggguaaaagccccc uugauuuugauuuucagugugaauacaaaccaugaaagugu   2751
            ** ***** *** **   ****** * *   ******      *   **    ****

Human       ggccucacgauccuucugaccuuuugguuuuaagcaggaggugucagaaaaguuaccac   4391
Mouse       ggccucacgauccuucugaccuuuugguuuuaagcaggaggugucagaaaaguuaccac   4046
Chicken     ggccucacgauccuucugacuuuuugguuuuaagcaggaggugucagaaaaguuaccac   3833
Fly         ggccuaucgauccuuuugguuuaaagaguuuuuaacaagaggugucagaaaaguuaccau   3344
C.elegans   ggccuaucgauccuuuuaauccugauuguuucagguaagaggugucagaaaaguuaccac   2924
Yeast       ggccuaucgauccuuuagucccucggaauuugaggcuagaggugccagaaaaguuaccac   2811
            *****  ********             ***       ****** **************

Human       agggauaacuggcuuguggcggccaagcguucauagcgacgucgcuuuuugauccuucga   4451
Mouse       agggauaacuggcuuguggcggccaagcguucauagcgacgucgcuuuuugauccuucga   4106
Chicken     agggauaacuggcuuguggcggccaagcguucauagcgacgucgcuuuuugauccuucga   3893
Fly         agggauaacuggcuuguggcggccaagcguucauagcgacgucgcuuuuugauccuucga   3404
C.elegans   agggauaacuggcuuguggcagccaagcguccauagcgacguugcuuuuugauccuucga   2984
Yeast       agggauaacuggcuuguggcagucaagcguucauagcgacauugcuuuuugauucuucga   2871
            ******************** * ******* ********* * ********** ******
```

```
Human       ugucggcucuuccuaucauugugaagcagaauucaccaagcguuggauuguucacccacu    4511
Mouse       ugucggcucuuccuaucauugugaagcagaauucaccaagcguuggauuguucacccacu    4166
Chicken     ugucggcucuuccuaucauugugaagcagaauucaccaagcguuggauuguucacccacu    3953
Fly         ugucggcucuuccuaucauugugaagcaaaauucaccaagcguuggauuguucacccaug    3464
C.elegans   ugucggcucuuccuaucauugcgaagcagaauucgccaagcguuggauuguucacccacu    3044
Yeast       ugucggcucuuccuaucauaccgaagcagaauucgguaagcguuggauuguucacccacu    2931
            *******************   ****** *****   ********************

Human       aauagggaacgugagcuggguuuagaccgucgugagacagguuaguuuuacccuacugau    4571
Mouse       aauagggaacgugagcugggauuagaccgucgugagacagguuaguuuuacccuacugau    4226
Chicken     aauagggaacgugagcuggguuuagaccgucgugagacagguuaguuuuacccuacugau    4013
Fly         -caagggaacgugagcuggguuuagaccgucgugagacagguuaguuuuacccuacuaau    3523
C.elegans   aauagggaacgugagcuggguuuagaccgucgugagacagguuaguuuuacccuacuguu    3104
Yeast       aauagggaacgugagcuggguuuagaccgucgugagacagguuaguuuuacccuacugau    2991
               ***************** ************************************  *

Human       gau---guguuguugccaugguaauccugcucaguacgagaggaaccgcagguucagaca    4628
Mouse       gau---guguuguugccaugguaauccugcucaguacgagaggaaccgcagguucagaca    4283
Chicken     gau---guguuguugcgcuaguaauccugcucaguacgagaggaaccgcagguucagaca    4070
Fly         gacaaaacguuguugcgacagcauuccugcguaguacgagaggaaccgcagguacggacc    3583
C.elegans   gac---uuguuauugcgaaaguaauccugcuuaguacgagaggaacagcggguucaaaca    3161
Yeast       gaa----uguuaccgcaauaguaauugaacuuaguacgagaggaacaguucauucggaua    3047
            **      ***   **    * * *    *  ************** *    * *  *

Human       uuugguguaugugcuuggcugaggagccaauggggcgaagcuaccaucugugggauuaug    4688
Mouse       uuugguguaugugcuuggcugaggagccaauggggcgaagcuaccaucugugggauuaug    4343
Chicken     uuugguguaugugcuuggcugaggagccacuggagcgaggcuaccaucugugggauuaug    4130
Fly         aauggcaca-auacuuguucgagcgaacagugguaugacgcuac-guccguuggauuaug    3641
C.elegans   uuugguucauagacuugaucgacagaucaauggucugaagcuaccauuugagagauuaua    3221
Yeast       auugguuuuugcggcugucugaucaggcauugccgcgaagcuaccauccgcuggauuaug    3107
              ***          **   **     ** **    ** *****  *  *   ******

Human       acugaacgccucuaagucagaaucccgcccaggcgg-aacgauacgg---cagcgccgc-    4743
Mouse       acugaacgccucuaagucagaaucccgcccaggcgg-aacgauacgg---cagcgccgaa    4399
Chicken     acugaacgccucuaagucagaaucccccuaaacgu-agcgauaccg---cagcgccga-     4185
Fly         ccugaacgccucuaaggucguauccgugcuggacugcaaugauaaau---aaggggc-aa    3697
C.elegans   acugaacgccucuaaguuagaaucucgccuugucaa-ggcgaaaauuucuugcuucc-c-    3278
Yeast       gcugaacgccucuaagucagaauccaugcuag--aa-cgcggugauuucuuugcuccac-    3163
             ***************   * ***    *           *               *

Human       ggagccucgguuggccucggauagccgguccccgccuguccccgccgg-cggccgccc      4802
Mouse       ggagccucgguuggccccggauagccgggucccgguccguccccgcucggcggggucccc    4459
Chicken     ggcgccucggugggcucgcgauagccggccgccgccc-----------------------    4222
Fly         uuugcauuguauggcuucuaa---accauuuaaaguuuauaauuu---------------    3739
C.elegans   ggugucgg--------------gaggcauc---ucuaucucguggca-------------    3308
Yeast       acaauauag-auggauacgaauaaggcguc---cuuguggcgucgcu-------------    3206

Human       ccccccuccacgcgccccgcgcgcgcgggagggcgcgugccccgccgcgcgccgggaccg    4862
Mouse       gcgucguccccgcg-------gc--ggcgcggggucuccccccgccgggcgucgggaccg    4510
Chicken     ---------------cccu------cg----------------------------ggcgg    4233
Fly         acuuuau-----------------aaacgacaauggauguga-------------ugcca    3769
C.elegans   ------------------------acacgagagcuuaugccc-------------uaugu    3331
Yeast       ------------------------gaaccauagcaggcuagc-------------aa-cg    3228

Human       ggguccggugcggagugcccuucguccugggaaacggggcg---cggccg-gaaaggcgg    4918
Mouse       ggguccggugcggagagccguucgucuugggaaacggggug---cggccg-gaaaggggg    4566
Chicken     gcggucggugcggagcgccgcucguggucgggaccggagcg---cggaca-gauguggcg    4289
Fly         auguaauuug--uaa----------cauaguaaauugggaggaucuucgaucaccugaug    3817
C.elegans   auggccuuggcgucgu--agugaauucugcg-------acg---cuu------------g    3367
Yeast       gugcacuuggcgga-------aaggccuugg-------gug---cuu------------g    3259
              *      *                 *            *   *              *
```

```
Human       ccgccccc-ucgcccgucacgc--accgcacguu--cgug-gggaaccuggcgcuaaacc    4972
Mouse       ccgcccuc-ucgcccgucacguugaacgcacguu--cgug-uggaaccuggcgcuagacc    4622
Chicken     ccg-ccuc-ucccccgccgcgu--accgcauguu--cgug-gggaacccggugcuaaauc    4342
Fly         ccgcgcuaguuacaua------uaaaagcauuauuuaauacaaugacaaagccuagaauc    3871
C.elegans   ccaacg-----ccagaucacucug-------guu--ca----augucggggcgcuaaauc    3409
Yeast       cuggcgaauugcaaugucauu----------------------uugcguggggauaaauc    3297
            *                                             *   *      * *

Human       auucguagacgaccugcu-ucugggucgggguuucguacguagcagagcagcucccuc--    5029
Mouse       auucguagacgaccugcu-ucugggucgggguuucguacguagcagagcagcucccuc--    4679
Chicken     auucguagacgaccugau-ucugggucgggguuucguacguagcagagcagcucccuc--    4399
Fly         aauuguaaacgacuuuug-uaacaggcaagguguuguaagugguugagcagcugccau--    3928
C.elegans   acuugcauacgacuuggucucuuggucaagguguuguauucaguagagcaguccuuuuau    3469
Yeast       auuuguauacgacuuaga-uguacaacgggguauuguaagcaguagaguagccuuguu--    3354
            * * * * ***** *    *      *  *** * ***    *  *** **

Human       gcugcgaucuauugaaagucagcccucgacacaaggguuuguc-------------    5072
Mouse       gcugcgaucuauugaaagucagcccucgacacaaggguuugucucugcgggcuuuc    4735
Chicken     gcugcgaucuauugagagucagcccucgacacaagcuuuuguc-------------    4442
Fly         acugcgauccacugaagcuuauccuuugcuugau-gauucga--------------    3969
C.elegans   acugcgaucuguugagacuauccuuug-auugaguuuuuug---------------    3509
Yeast       guuacgaucugcugagauuaagccuuu-guugucugauuugu--------------    3395
              * *****   ***   *   *  *           ** *
```
